# Patterns of aDNA Damage Through Time and Environments – lessons from herbarium specimens

**DOI:** 10.1101/2025.10.26.684600

**Authors:** Stefano Porrelli, Alice Fornasiero, Hong Phuong Le, Wenzhe Yin, Maria Navarrete Rodriguez, Nahed Mohammed, Axel Himmelbach, Andrew C. Clarke, Nils Stein, Paul J. Kersey, Rod A. Wing, Rafal M. Gutaker

## Abstract

Herbarium collections are a vast but underutilized resource for ancient DNA research, containing over 400 million specimens with detailed metadata and spanning centuries of global biodiversity. Understanding patterns of DNA preservation in natural collections is crucial for optimizing ancient DNA studies and informing future curation practices. We analysed genomic data for 573 herbarium specimens from six plant species from the genera *Hordeum* and *Oryza* collected from the Americas and Eurasia over 220 years. Using standardized laboratory protocols and shotgun sequencing, we quantified DNA degradation and elucidated factors that accelerate it. We find significant age-dependent DNA fragmentation rates, indicating temporal degradation processes not detected in prehistoric samples. In our analysis, DNA decay rates in herbarium specimens were almost eight times faster than in moa bones, reflecting fundamental differences in tissue composition and preservation environments. Environmental conditions at the time of specimen collection emerged as the major determinants of post-mortem damage rates, with the interaction term between temperature and genus being the dominant driver of cytosine deamination. We find no effect of sample storage on DNA damage and degradation. These findings provide insights into how climatic origin, preservation environment, taxonomic identity and age influence DNA preservation while highlighting opportunities for improving institutional preservation practices. Due to standardised preservation conditions, museum collections can provide better insights into DNA damage and degradation over time than archaeological and paleontological samples.

## Introduction

Understanding preservation of DNA from old biological samples, commonly referred to as ancient DNA (aDNA), has been at the core of the genomic revolution in archaeological research [1,2]. There is a consensus among researchers that aDNA is defined by its degradation and not by its age [3,4], though the two are correlated [5]. The two most prominent *post-mortem* reactions associated with DNA damage and degradation are deamination and depurination, which occur spontaneously in the absence of enzymatic repair machinery [6,7]. Deamination of cytosines into uracils leads to a characteristic pattern of ‘C>T’ misincorporations, with increased frequency at fragment termini [8], whilst depurination causes “nicks” (breakage) in the phosphodiester bonds and subsequent hydrolysis of DNA backbone, resulting in DNA fragmentation towards very small molecules (∼30-100 bp). Both patterns are common in historical, archaeological and sedimentary samples [2,5,9] and make bioinformatic processing and downstream analyses challenging [10]. Although 5’ C>T misincorporations (and the complementary 3’ G>A misincorporations in double-stranded libraries [3]) can be used to authenticate genuine aDNA sequences, they can bias variant calling and phylogenetic analyses, potentially leading to incorrect inferences [3]. Similarly, extensive DNA fragmentation reduces mapping efficiency and increases the likelihood of spurious alignments [11,12]. These challenges are particularly acute for ancient samples from hot and humid environments, where accelerated degradation processes further compromise DNA integrity [13,14]. As a result, tools and approaches have been developed to improve quality control, reads processing, mapping and downstream analyses in a quest to utilize even highly degraded samples [15–17]. In addition to chemical degradation, DNA can also be altered through biological processes such as microbial colonization. In effect, the *bona fide* DNA from the target species can be depleted and replaced with post-mortem microbial DNA [18].

While the vast majority of published aDNA research focuses on human skeletal remains [19,20], there are an increasing number of studies in other mammals [21], arthropods [9,22], and in plants [23–25]. Archaeobotanical materials, primarily seeds, are often used as sources of prehistoric DNA. While not older than 500 years, herbarium specimens (figure 1) number over 400 million [26], have solid species identification and, frequently, collection metadata. They comprise a vast but underutilised resource for studying evolutionary and ecological changes in the last couple of centuries [27–30]. The leaf tissue is most often the target in DNA isolation protocols but does not possess the same structural isolation from the environment as bones or seeds. This is the most likely reason why DNA in herbarium samples decays at a rate roughly six times faster than in ancient Moa bones [25], but also more than twice as slow when compared to dry-pinned arthropod museum specimens [9]. On the other hand, herbarium specimens are commonly dried upon collection and are generally stored in standardized conditions that are favourable for DNA preservation, such as stable temperatures and low air humidity. Availability of materials, good geolocation information and relatively stable preservation condition *ex situ* make herbarium specimens a perfect system to study the effects of the environment at the point of collection on the subsequent preservation of DNA in biological samples.

**Figure 1:**
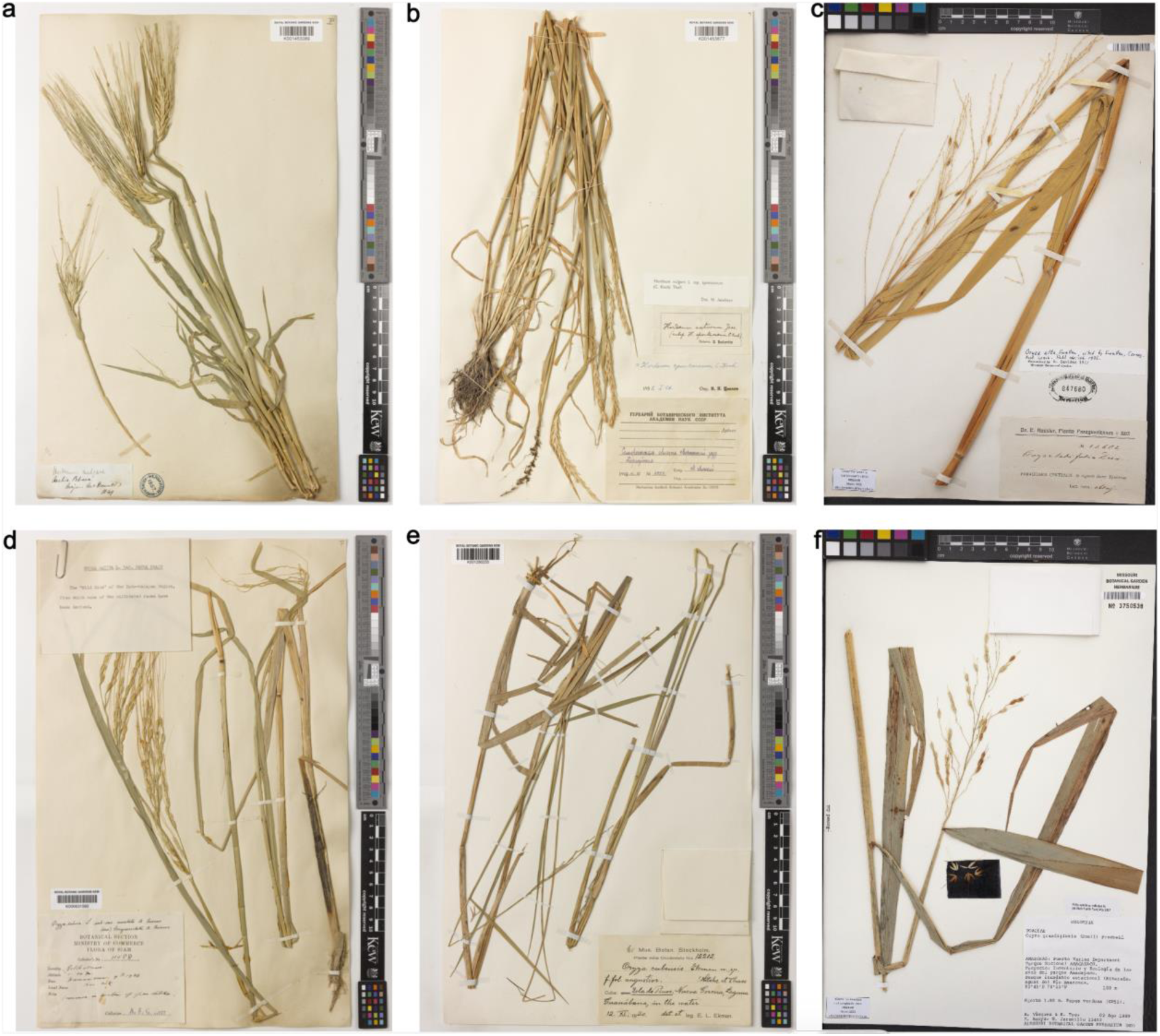
Representative herbarium specimens for the six species analysed in this study. Photos courtesy of Royal Botanic Gardens, Kew (RBGK) and Missouri Botanic Gardens (MBG). (a) *Hordeum vulgare* sample HV0080, collected in 1849 (RBGK); (b) *Hordeum spontaneum* sample HS0156, collected in 1909 (RBGK); (c) *Oryza alta* sample HRG0858, collected in 1913 (MBG); (d) *Oryza rufipogon* sample HRG0849, collected in 1926 (RBGK); (e) *Oryza latifolia* sample HRG0287, collected in 1920 (RBGK); (f) *Oryza grandiglumis* sample HRG0580, collected in 1989 (MBG).

One of the major challenges in studies trying to investigate the dynamics of DNA damage and fragmentation is lack of large datasets with a global distribution and consistent sampling and laboratory processing procedures. Previous studies were limited to one or two species representing narrow geographical range [13,25]. Here, we present new sequencing data for a total of 573 herbarium samples from six plant species, spanning the Americas and Eurasia, processed with the same laboratory protocol in dedicated aDNA laboratory facilities, and sequenced using a whole-genome shotgun approach. Our main aim is to understand which environmental factors influence the rates of DNA damage and decay. The outcome of this investigation brings new insights into the fundamental processes of DNA degradation that can inform future conservation of museum specimens, as well as help researchers to prioritize materials for their studies.

## Methods

### Herbarium samples

Historical herbarium samples (*N* = 573) were obtained from seven herbaria in Europe and North America (table 1): Royal Botanic Gardens, Kew (UK), Smithsonian Institute Herbarium (US), Missouri Botanic Gardens (US), New York Botanic Gardens (US), National History Museum (France), Harvard University Herbarium (US), and National History Museum (UK). Passport information for each sample is included in the supplementary material (supplementary table S1). Although we aimed at representing the majority of environments for each species in our dataset, some parts of distribution might have been overlooked due to unavailability of samples in our collections. The *Hordeum* samples were predominantly collected from temperate Europe to semi-arid regions of the Middle East, with highest counts from countries such as Iraq, Israel and Iran. The *Oryza* samples were predominantly collected from tropical and subtropical regions of the Americas and Southeast Asia, particularly Brazil, Costa Rica, Colombia, Mexico and Thailand (figure 2).

**Figure 2:**
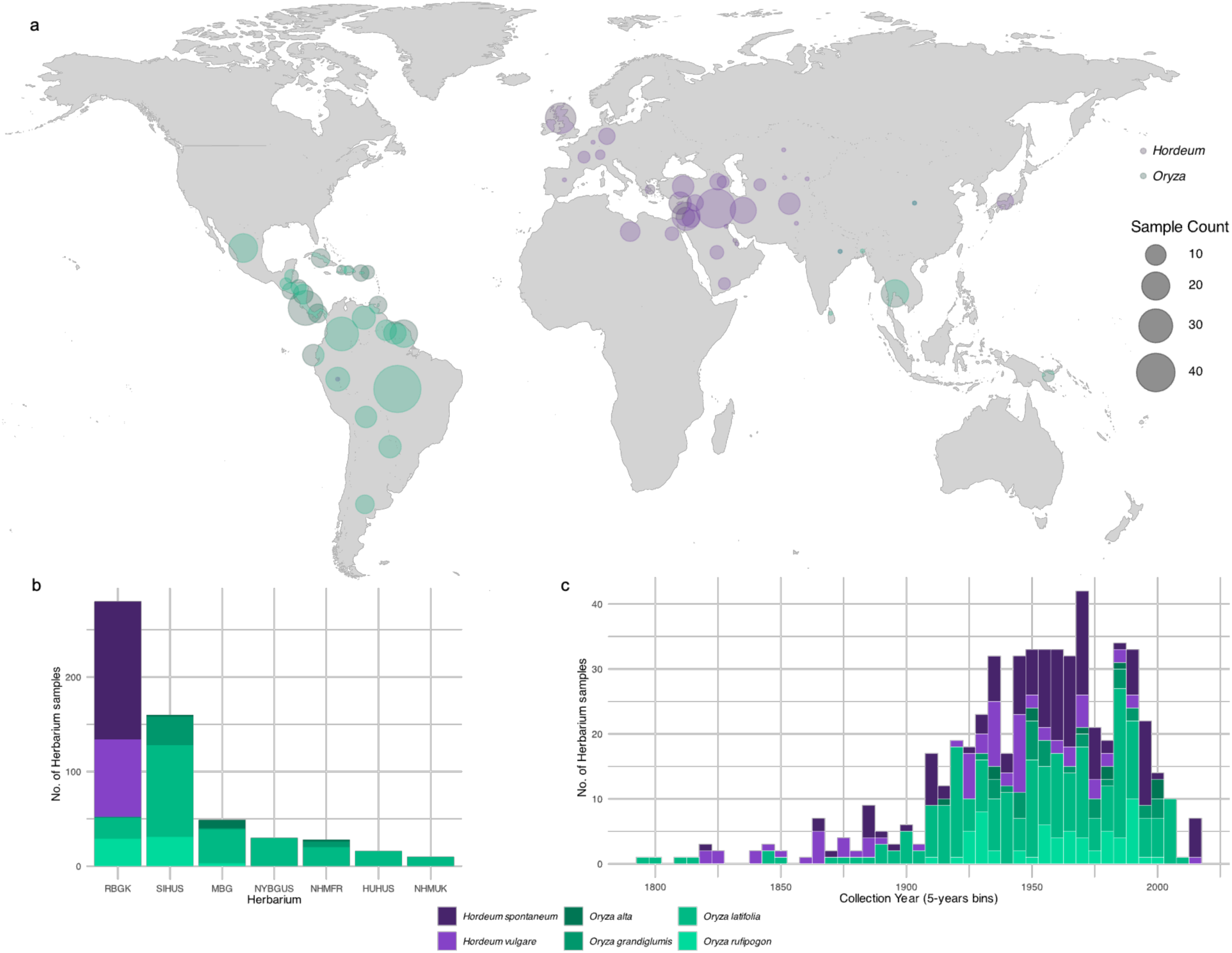
Overview of the herbarium samples of barley (*Hordeum*) and rice (*Oryza*) species: (a) geographical distribution based on country of collection; (b) source of herbarium of *Hordeum* and *Oryza* species (RBGK: Royal Botanic Gardens Kew, UK; SIHUS: Smithsonian Institute Herbarium, US; MBG: Missouri Botanic Gardens, US; NYBGUS: New York Botanic Gardens, US; NHMFR: National History Museum, France; HUHUS: Harvard University Herbarium, US; NHMUK: National History Museum, UK); (c) temporal distribution for each species.

**Table 1:**
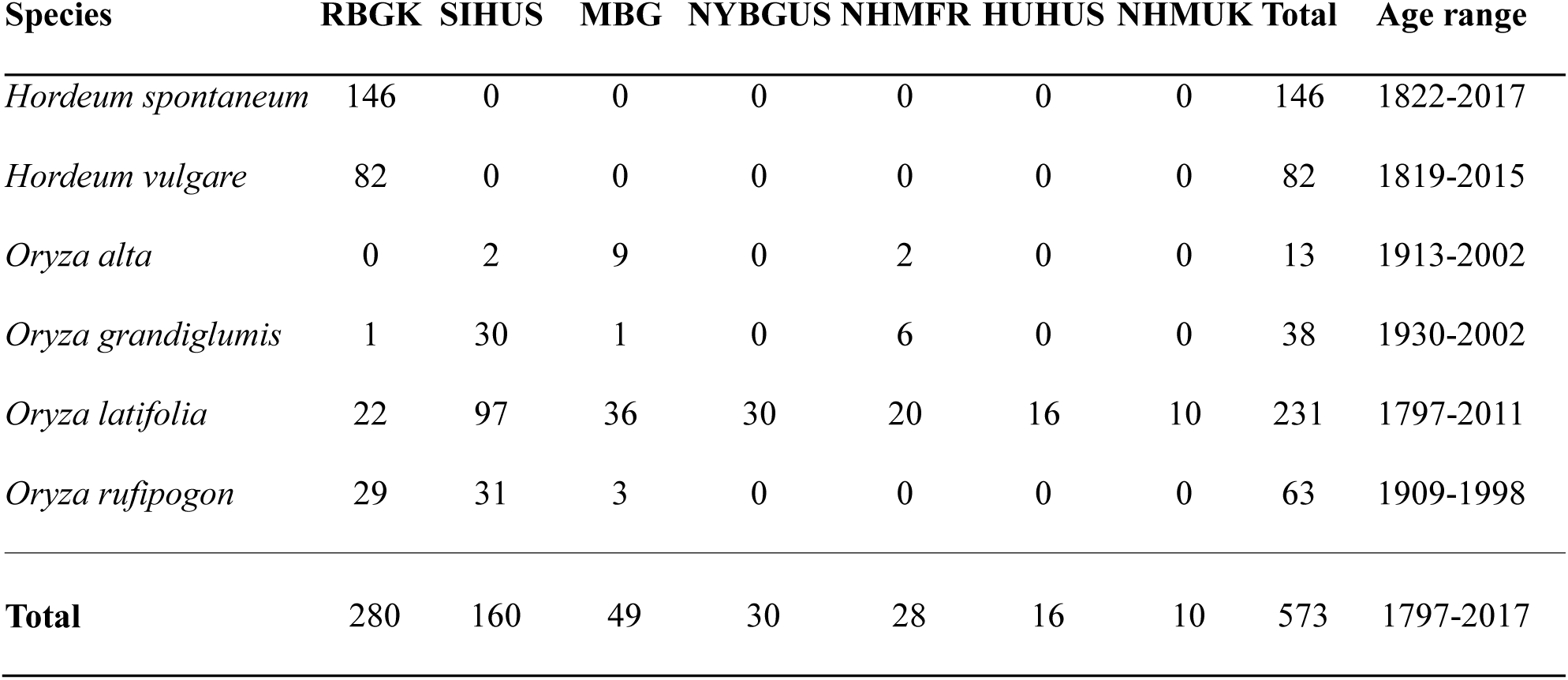
Number of historical herbarium samples (*N* = 573) included in this study collected from seven herbaria. RBGK: Royal Botanic Gardens Kew, UK; SIHUS: Smithsonian Institute Herbarium, US; MBG: Missouri Botanic Gardens, US; NYBGUS: New York Botanic Gardens, US; NHMFR: National History Museum, France; HUHUS: Harvard University Herbarium, US; NHMUK: National History Museum, UK.

| Species | RBGK | SIHUS | MBG | NYBGUS | NHMFR | HUHUS | NHMUK | Total | Age range |
| --- | --- | --- | --- | --- | --- | --- | --- | --- | --- |
| *Hordeum spontaneum* | 146 | 0 | 0 | 0 | 0 | 0 | 0 | 146 | 1822-2017 |
| *Hordeum vulgare* | 82 | 0 | 0 | 0 | 0 | 0 | 0 | 82 | 1819-2015 |
| *Oryza alta* | 0 | 2 | 9 | 0 | 2 | 0 | 0 | 13 | 1913-2002 |
| *Oryza grandiglumis* | 1 | 30 | 1 | 0 | 6 | 0 | 0 | 38 | 1930-2002 |
| *Oryza latifolia* | 22 | 97 | 36 | 30 | 20 | 16 | 10 | 231 | 1797-2011 |
| *Oryza rufipogon* | 29 | 31 | 3 | 0 | 0 | 0 | 0 | 63 | 1909-1998 |
| **Total** | 280 | 160 | 49 | 30 | 28 | 16 | 10 | 573 | 1797-2017 |

### DNA extraction, library preparation and low throughput sequencing

Laboratory steps for DNA extraction and library preparation for all 228 *Hordeum* (146 *H. spontaneum* and 82 *H. vulgare)* and 58 *Oryza* (30 *O. latifolia,* 18 *O. rufipogon* and 10 *O. grandiglumis*) (supplementary table S1) samples were carried out in a clean room facility at Royal Botanic Gardens, Kew (RBGK, UK) following previously published best practices and aDNA protocol described in [31] with adjusted volumes. A further 287 *Oryza* samples (201 *O. latifolia*, 45 *O. rufipogon*, 28 *O. grandiglumis* and 13 *O. alta*) (supplementary table S1) were processed in a clean room facility at the Ancient and Environmental DNA Laboratory (ÆDNA) at the University of Nottingham (UoN, UK), following the same protocol and volume adjustments. Following best practices for aDNA research [3,31], negative controls were processed alongside samples to monitor for environmental and reagent contamination. These included extraction blanks and library preparation controls.

Leaf tissue samples (5-10 mg) were grinded in 2mL PowerBead metal tubes (QIAGEN, 13117-50) with a bead mill homogeniser (Precellys Evolution, P002511-PEVT0-A.0). 1mL of PTB-based mix was added to the homogenized samples, which were then incubated on a rotor overnight at 37°C. Genomic DNA (gDNA) was isolated and half the volume (0.5 mL) was purified using the DNeasy Plant Mini Kit (QIAGEN, 69106) with modifications described in [31], and quantified with Quantus™ fluorometer (Promega, E6150). A subset of the gDNA isolates (*N*=40, 10 *H. spontaneum*, 10 *H. vulgare*, 10 *O. rufipogon*, and 10 *O. latifolia*; supplementary table S2) with concentrations > 5 ng/μL were selected for measurements of gDNA fragment size distributions with capillary electrophoresis. These samples were analysed on a TapeStation 4200 system (Agilent, G2991BA) using a D1000 ScreenTape (Agilent, 5067-5582).

For all 573 samples, double stranded and doubled indexed genomic libraries were constructed by blunt-end DNA ligation [32,33] following [31]. Briefly, genomic DNA was blunt-end repaired, and universal Illumina double-stranded adapters were ligated. This was followed by fill-in of adapters, indexing with unique combinatorial Illumina indexes, and amplification.

Between each step, samples were purified with the MiniElute® PCR purification kit (QIAGEN, 28006). To quantify the excess of 5’ C>T misincorporations at reads termini caused by spontaneous deamination of cytosines [34], and to validate aDNA authenticity [25,35], we did not perform enzymatic removal of aDNA-associated DNA misincorporation [36]. Indexed and amplified libraries were quantified with Quantus™ fluorometer (Promega, E6150). Fragment size of each library were estimated with a 4200 TapeStation System (Agilent, G2991BA) using a D1000 ScreenTape (Agilent, 5067-5582).

Libraries were pooled equimolarly and sequenced in paired-end mode. All 228 *Hordeum* libraries processed at RBGK were sequenced on a NovaSeq 6000 or NovaSeq X+ device (2 x 62 bp) according to manufacturer’s instructions (Illumina, San Diego, CA, USA) at the Leibniz Institute of Plant Genetics and Crop Plant Research (IPK, Gatersleben, Germany). The 58 *Oryza* libraries processed at RBGK were sequenced at Macrogen (Macrogen, Europe), on an Illumina Novaseq X+ system (2 x 150 bp). The 287 *Oryza* samples processed at the Ancient and Environmental DNA Laboratory (ÆDNA) at the University of Nottingham (UoN, UK) were sequenced on an Illumina Miseq platform (2 x 150 bp) according to manufacturer’s instructions (Illumina, San Diego, CA, USA) at the Deep Seq facility at UoN.

### Bioinformatic screening and validation of aDNA authenticity

Bioinformatics protocols for read processing, quality assessment, screening and authentication of aDNA-derived libraries follow [31]. Demultiplexed raw Illumina paired-end reads were adapter trimmed and merged with AdapterRemoval v2.3.4 [37] with a minimum overlap of 11 bases, and quality-checked with FastQC v0.11.8 (http://www.bioinformatics.bbsrc.ac.uk/projects/fastqc). Reference genome assemblies were retrieved from the National Center for Biotechnology Information (NCBI) database (table 2) and indexed with BWA v0.7.19 [38] and SAMtools v1.21 [39]. For each species, quality- and adapter-trimmed merged reads were mapped to their respective reference genome with BWA-aln using aDNA-specific parameters (“-l 1024”). PCR optical duplicate reads were removed with DeDup [17]. Base frequencies at DNA break points and nucleotide misincorporations were estimated for each library using MapDamage2 v2.0.6 [40]. Mapping statistics of the additional 287 *Oryza* samples were obtained by mapping sequencing reads to the following reference genomes: *O. alta* (PRJNA1039467), *O. grandiglumis* (PRJNA737282), and *O. latifolia* (PRJNA737486) [41]. The *O. rufipogon* samples originally collected in the American continent were aligned to the *O. glumipatula* genome reference (PRJNA48429). Negative laboratory controls (extraction and library blanks) were included in the bioinformatic screening to monitor/assess possible contamination, but they were excluded from further analyses.

**Table 2:** Reference genome assemblies used in this study with associated BioProject, BioSample and Accession IDs.

| Reference | BioProject | BioSample | Assembly Accession |
| --- | --- | --- | --- |
| *Hordeum spontaneum* | PRJEB57567 | SAMEA112465237 | GCA_949783245.1 |
| *Hordeum vulgare* | PRJEB40589 | SAMEA7384724 | GCF_904849725.1 |
| *Oryza alta* | PRJNA1039467 | SAMN38217704 | GCA_047899615.1 |
| *Oryza grandiglumis* | PRJNA737282 | SAMN19687255 | GCA_048188845.1 |
| *Oryza latifolia* | PRJNA737486 | SAMN19696891 | GCA_048174585.1 |
| *Oryza rufipogon* | PRJNA1029807 | SAMN40302812 | GCA_037997075.1 |
| *Oryza glumipatula* | PRJNA48429 | SAMN02981440 | GCA_000576495.2 |

### Ancient DNA damage metrics and regression analyses

Four metrics were selected to quantify patterns of aDNA damage: (i) the proportion of endogenous DNA content, (ii) the fragment length distribution, (iii) the damage fraction per site (λ), and (iv) the frequencies of 5’ C>T misincorporations at the first base. The four metrics were analysed in linear models as a function of collection year and sample age using the ‘*lm*’ function in R [42]. Model assumptions were verified using standard diagnostic procedures. For all linear regressions, we examined residual plots to assess the assumptions of normality and homoscedasticity.

#### Endogenous content

The proportion of endogenous DNA was calculated for each sample as the fraction of quality-and adapter-trimmed merged reads that mapped to their respective reference genome divided by the total number of reads retained after adapter-trimming and quality filtering. Only primary alignments were considered, with mapped reads representing unique mapping decision for each query sequence. Since this metric can be impacted by increasing evolutionary distance between target and reference [9], we used the most closely related reference genome available for each of the species included in our dataset (table 2). Endogenous content was calculated with SAMtools “*flagstat*” tool [39] and plotted as a function of collection year.

#### Fragment length

DNA fragmentation can be quantified by agarose gel electrophoresis, automated electrophoresis or by the *in-silico* generation of fragments by merging overlapping paired reads of Illumina libraries [43,44]. To validate that the fragment sizes of the merged reads reflects the original molecule length, we examined the relationship between the fragment size distribution of isolated gDNA and that of the amplified libraries for a subset of the samples (*N=* 40; 10 *H. spontaneum*, 10 *H. vulgare*, 10 *O. rufipogon*, and 10 *O. latifolia*; supplementary table S2) using TapeStation profiles. Library and gDNA peak size measurements from the TapeStation profiles were compared with each other and with bioinformatically-derived median fragment lengths of merged reads to assess the concordance between direct physical measurements and computational estimates. In addition, we further analysed the correlation between peak sizes of the gDNA isolates and age of the sample. Having established a relationship between fragment size distributions of gDNA and bioinformatically-derived merged reads for a representative subset (*N*=40), we examined the relationship between fragment size of merged reads and age for the whole dataset (*N=* 573). We fitted the fragment length distribution of the mapped merged reads to a lognormal distribution using the ‘fitdistr’ function from the MASS package [45] in R. We used the mean of this distribution (log-mean) to summarise fragment length for each library and carried out regressions on the relationship between the log-mean of fragment lengths and collection year. As suggested in [25], we used the median plotted on a log-scaled *y-*axis for visualisation, as the latter is more intuitive to understand than the log-mean.

#### Damage fraction per site (λ) and DNA decay rate (k)

The damage fraction per site (λ) was calculated for each sample using previously described methods [13]. We fitted an exponential decay model to the empirical fragment length distribution of the mapped reads and derived λ as the negative of the slope coefficient of the linear regression. To identify samples deviating from the assumption of exponential decay of fragment length, we assessed goodness of fit and statistical significance on a per-sample basis. Model fit was considered satisfactory when *R^2^* > 0.95 and *p <* 0.05.

Having determined the damage fraction per site (λ), we calculated genus-specific DNA decay rates (*k*) by plotting λ as a function of sample age and extrapolating *k* from the slope of the linear regression, according to the linear relationship *k* = *λ*⁄*age* [46]. We also calculated the overall DNA decay rate (*k*) of all herbarium samples irrespective of genus.

#### Nucleotide misincorporations

The frequency of 5’ C>T misincorporations at first position was used as a proxy for 5’ damage. Although the depth of sequencing needed for reliable estimation of C>T frequencies is sample-dependent, an order of magnitude of thousands of reads is considered sufficient [47]. Therefore, only samples with >5,000 merged paired reads were retained for further analyses. Furthermore, the exponential increase of C>T misincorporations at reads termini is routinely used as a metric to validate authenticity of aDNA [5]. We scored this pattern for each sample by fitting an exponential model to the C>T frequencies for the first 20 bases at 5’ terminus. We evaluated the goodness of fit with a one-sided *t-*test on a per-sample basis, as described in [48], and only retained samples that fitted the exponential model (*R^2^* > 0.5, *p <* 0.05). Finally, we carried out a regression analysis of 5’ C>T damage as a function of sample age.

For comparison, and to demonstrate that both 5’ and 3’ ends in double-stranded libraries show concordant deamination patterns, we also analysed the complementary 3’ G>A misincorporations. To demonstrate the patterns observed are specific to deamination, we computed the average frequency of non-deamination substitutions for all samples following the approach described in [12], and plotted this as a function of sample age. To account for increased rates of baseline substitutions arising from evolutionary divergence between samples and reference genomes, we calculated mean divergence from reference on a species-basis. We also determined divergence-corrected deamination rates by subtracting baseline substitution frequencies from total deamination frequencies and compared the results against the non-corrected deamination rates.

### Analysis of covariance

To investigate differences in aDNA damage patterns between *Hordeum* and *Oryza*, we performed Analysis of Covariance (ANCOVA) using the ‘anova’ function in R. For each aDNA damage metric, sample age was used as the covariate and the genus as the factor using the model “*y* ∼ *covariate* x *factor”,* which also tested for possible interactions between sample age and genus (i.e., differences in the slope of regression are dependent on genus). We tested this model against a model of type “*y* ∼ *covariate* + *factor”* to assess whether the removal of the interaction influenced model fit. When the interaction was not significant (*p* > 0.05) we accepted the simpler model without interaction and concluded that regression slopes did not differ between genera, though intercepts might [25].

### Climate analyses

We obtained climate data from the CHELSA V2.1 climate dataset [49] to investigate the relationship between specimen preservation conditions, aDNA damage patterns, and climatic variables. Four primary bioclimatic variables were extracted for each sample location based on latitude and longitude metadata: (i) annual mean temperature (bio1), (ii) temperature seasonality (bio4), (iii) annual precipitation (bio12), and (iv) precipitation seasonality (bio15), henceforth referred as “annual climate”. Additionally, we retrieved monthly temperature (tas_01 - tas_12) and monthly precipitation (pr_01 - pr_12) means for year 1981-2010 (CHELSA V2.1) corresponding to each sample’s recorded month of collection and geographical location. We used this data to infer climatic conditions at the time of specimen collection, henceforth referred as “collection climate”.

To quantify the unique and shared contributions of multiple explanatory variables to the total variance in aDNA damage metrics, we performed a variance partitioning analysis using the ‘varpart’ function implemented in the VEGAN package [50]. For each aDNA metric, we applied a “collection climate” model, where monthly climatic variables (temperature and precipitation) were assigned to samples based on their recorded location and month of collection, and an “annual climate” model, where annual mean temperature, mean precipitation and their seasonality were assigned to samples based on their geographical location. The “collection climate” model aimed at capturing DNA damage occurring during the initial post-collection period (field handling, drying, and early preservation), when specimens are exposed to ambient environmental conditions, rather than during subsequent long-term storage under standardised herbarium conditions [2,25,51]. The “annual climate” model aimed at capturing the general climatic regime of the collection locality, thus providing a comparison to assess whether month-specific climate data improves explanatory power over annual averages. Annual climate variables may also better represent cumulative environmental exposure if pre-collection conditions (e.g., growing season climate) influence tissue properties relevant to DNA preservation.

As a possible confounding effect, we also included genus as a variable, as well as herbarium, which could be interpreted as different long-term institutional storage conditions. Since ‘varpart’ function accommodates only up to four variables, for some analyses, temperature and precipitation were merged into single climatic variable. For each response (endogenous DNA content, fragment size, lambda and 5’ C>T damage), statistical significance of individual fractions and combinations were tested using redundancy analysis with the ‘rda’ function implemented in VEGAN, with 999 permutation followed by ANOVA.

### Divergence-corrected nucleotide misincorporations analyses

To assess whether reads-reference divergence influenced our main findings, we repeated all primary analyses using divergence-corrected deamination frequencies. For each sample, corrected 5’ C>T frequencies were calculated by subtracting the mean baseline substitution rates from the observed 5’ C>T frequencies. Specifically, we re-ran the regression analysis of corrected deamination frequencies against sample age, the variance partitioning analyses to quantify the relative contributions of temperature, precipitation, age, and genus to variation in corrected deamination, and the regressions between corrected deamination and temperature.

## Results

A total of 573 herbarium specimens were sequenced, generating libraries sequenced to variable read counts and depth (supplementary table S1). *Hordeum* samples (*N =* 228) processed at the Royal Botanic Gardens, Kew (RBGK) generated an average of 12.08 ± 5.94 million reads per library, with 86.2 ± 24.3% of reads mapping to the relevant reference genome. *Oryza* samples (*N=* 58) processed at the RBGK yielded an average of 1.41 ± 0.93 million reads per library, with 86.5 ± 14.6% of reads mapping to the relevant refence genome. The remaining *Oryza* samples processed at the University of Nottingham (*N=* 287) generated an average of 0.06 ± 0.09 million reads per library, with 78.0 ± 26.2% of reads mapping to the relevant reference genome.

Baseline substitution rates, calculated as the mean frequency non-deamination substitution types, served as a proxy for evolutionary distance between samples and their respective reference genomes. Overall, sample-reference divergence rates were low across all species, ranging from 0.53% to 0.88% (supplementary figure S1). *Hordeum* species showed the lowest divergence from their respective reference genome, with *H. vulgare* at 0.53% ± 0.52% and *H. spontaneum* at 0.57% ± 0.27%. Among *Oryza* species, *O. grandiglumis* showed the lowest divergence (0.68% ± 0.16%), followed *by O. rufipogon* (0.76% ± 0.31%), *O. latifolia* (0.80% ± 0.34%), and *O. alta* (0.88% ± 0.08%).

Damage pattern analysis revealed characteristic aDNA signatures. *Hordeum* samples exhibited an average 5’ C>T misincorporation frequency of 1.09 ± 0.44% at the first base of sequenced molecule, whilst *Oryza* samples exhibited an average of 2.13 ± 0.70%. These damage signatures, combined with fragment length distributions, were used to authenticate samples and filter out those inconsistent with genuine aDNA characteristics. Our filtering strategy identified samples deviating from the assumption of exponential decay of fragment length, as well as samples that did not display an exponential increase of C>T misincorporations at read termini diagnostic of aDNA. In total, 117 (20%) samples were removed, leaving a final dataset containing 456 samples.

### Regression analyses

#### Endogenous fraction

We analysed the proportion of reads mapping to the reference genome as a proxy for endogenous DNA content. The regression analyses revealed no statistically significant relationship between the proportion of endogenous DNA and the sample collection year in *Hordeum* (*R^2^ =* 0.003, *p* = 0.451, *N* = 211), but a very weak yet significant relationship was observed in *Oryza* (*R^2^ =* 0.04, *p* = 0.00167, *N*= 245; figure 3a). As we aimed at investigating the effect of genus on the rates of aDNA damage, we also carried the regression for all samples irrespective of genera, which provided comparable results (*R^2^ =* 0.012, *p* = 0.0215, *N*= 456, supplementary figure S2a).

**Figure 3:**
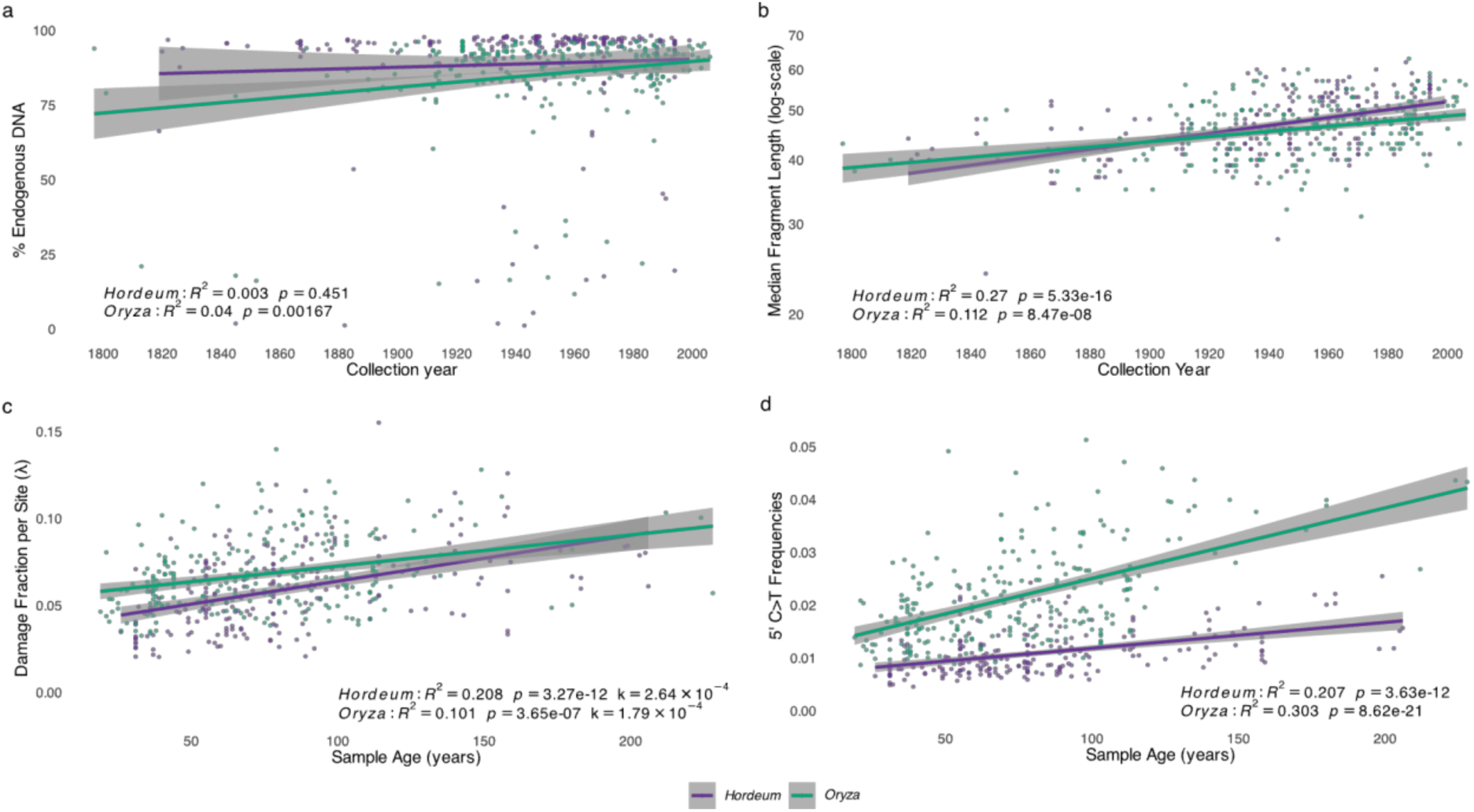
Regression analyses of aDNA damage metrics for *Hordeum* and *Oryza*: (a) Endogenous DNA content as a function of collection year. (b) Median fragment length of merged reads as a function of collection year, with log-scaled y-axis to show exponential relationship. (c) Damage fraction per site (λ) as a function of sample age, with the slope of regression corresponding to the DNA decay rate per base per year for *Hordeum* (*k = 2.64 × 10^-4^*) and *Oryza* (*k = 1.79 × 10^-4^*). (d) Frequencies of C>T misincorporations at first base (5’ -end) as a function of sample age. Insets show regression statistics for each aDNA damage metric for each genus.

#### Fragment length

We measured DNA fragment size using two complementary approaches. First, we validated our bioinformatic estimates of fragment size distributions using TapeStation profiles of both gDNA and amplified libraries for a subset of samples (*N=* 40). Peak size of amplified libraries strongly correlated with the bioinformatically-derived median fragment size of the merged library reads (*R^2^ =* 0.614, *p* =2.22 × 10^-9^, supplementary figure S3a), compared to the weaker correlation between the TapeStation peaks of libraries and gDNA origin, (*R^2^ =* 0.129, *p* =0.029, supplementary figure S3b). A significant, albeit weaker correlation was also observed between the gDNA peak size and the median fragment size of merged reads (*R^2^ =* 0.287, *p* =6.35 × 10^-4^, supplementary figure S3c), indicating that merging of overlapping reads of short insert libraries reflects, at least in part, the original molecule length [32]. The weaker correlation was expected, as library preparation and sequencing involve processing and purification steps that can impact the fragment size distribution. Furthermore, we observed a strong and significant relationship between gDNA peak size and collection year (*R^2^ =* 0.61, *p* =3.04 × 10^-8^, *N*= 40, figure 4), but only after removing the two *Hordeum* outliers HV0061 and HV0081, which were particularly old (collection years: 1842 and 1867, respectively) but showed gDNA size distribution patterns inconsistent with their age, possibly due to modern contamination of exogenous DNA. With the inclusion of these two outliers, we still observed a significant albeit weaker relationship (*R^2^ =* 0.263, *p* =1.19 × 10^-3^, supplementary figure S4).

**Figure 4:**
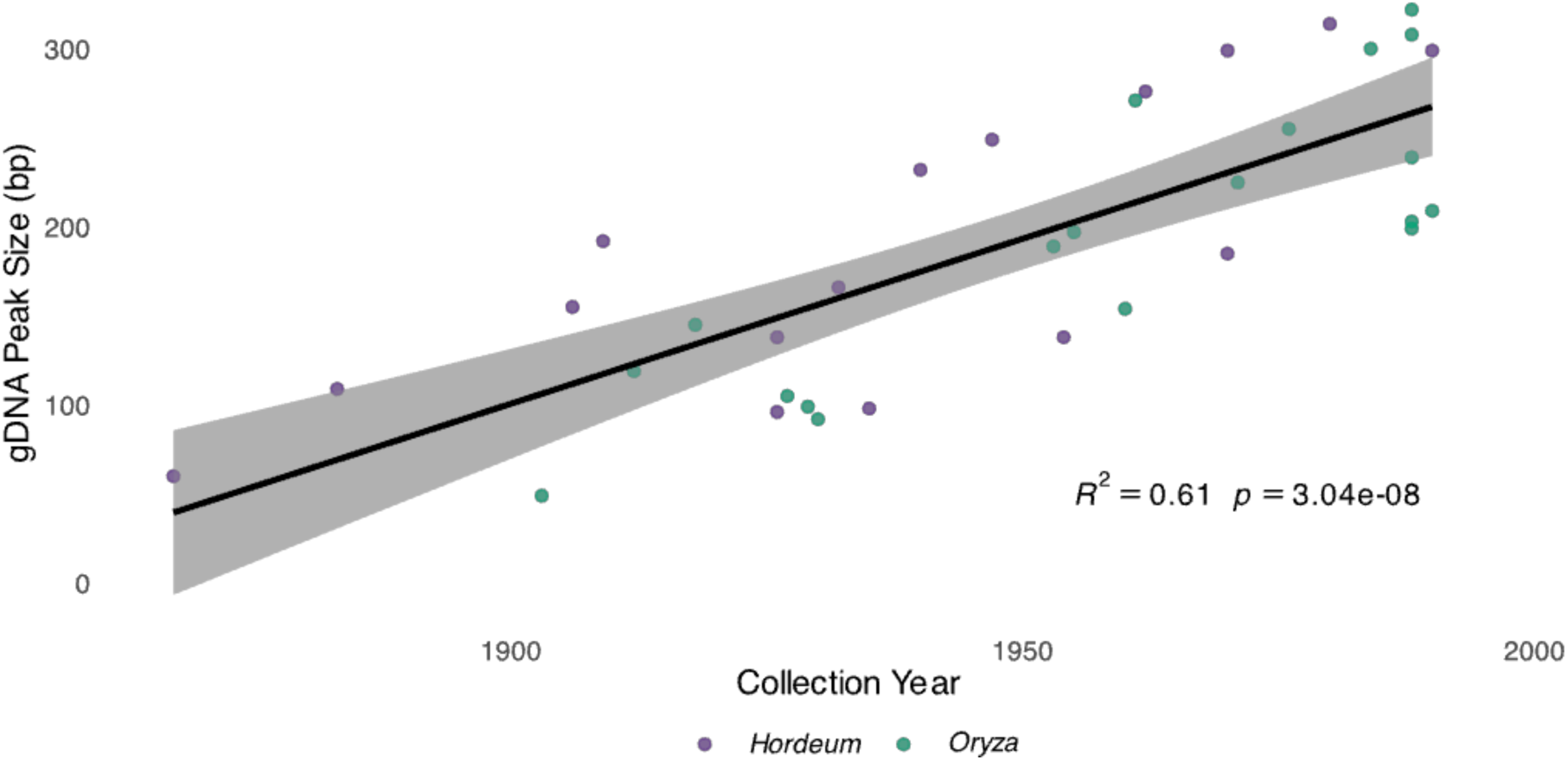
Regression between peaks of genomic DNA (gDNA) fragment size distribution obtained from high resolution fractionation electrophoresis (TapeStation) and collection year for a subset of samples (*N* = 40; 10 *H. spontaneum*, 10 *H. vulgare*, 10 *O. rufipogon*, and 10 *O. latifolia*;) after outlier removal. Inset shows regression statistics.

Second, we extended this analysis to the full dataset and examined the relationship between bioinformatically-derived fragment size and age. We observed a statistically significant relationship between the log-mean fragment length and the sample collection year for both genera (figure 3b), with a stronger relationship for *Hordeum* (*R^2^ =* 0.27, *p* =5.33 × 10^-16^, *N*= 211) than *Oryza* (*R^2^ =* 0.112, *p* = 8.47 × 10^-8^, *N*= 245). A statistically significant relationship was also observed when analysing all samples irrespective of genera (*R^2^ =* 0.171, *p* = 3.16 × 10^-20^, *N*= 456; supplementary figure S2b).

#### Damage fraction per site (λ) and DNA decay rate (k)

The slope of log-transformed exponential decline of fragment length frequencies in aDNA (λ) describes the probability of bond breaking in DNA backbone [46]. We estimated the DNA decay rate per year (*k*) for *Hordeum* and *Oryza* from the slope of the linear relationship between λ and sample age (figure 3c). We observed a per nucleotide decay rate of *k=* 2.64 × 10^-4^ per year for *Hordeum* (*R^2^ =* 0.208, *p* =3.27 × 10^-12^, *N*= 211), which was 1.5 times faster than the decay rate of *Oryza* of *k=* 1.79 × 10^-4^ per year (*R^2^ =* 0.101, *p* = 3.65 × 10^-7^, *N*= 245). The overall decay rate for all herbarium samples was *k*= 2.08 × 10^-4^ per year (*R^2^ =* 0.129, *p* = 2.52 × 10^-15^, *N*= 456, supplementary figure S2c), which is slightly faster than the decay rate of *k*= 1.66 × 10^-4^ per year observed in *Arabidopsis* and *Solanum* herbarium specimens [25], approximately 2.2 times slower than the *k* = 4.6 × 10⁻⁴ per year observed in dry-pinned arthropod museum specimens [9], and nearly eight times faster than the rate of *k*= 2.71 × 10^-5^ observed for ancient moa bones [13].

#### Nucleotide misincorporations

Deamination frequencies at 5’ (C>T) and 3’(G>A) were highly correlated (*R^2^ =* 0.951, *p* = 7.14 × 10^-161^, *N*= 245 for *Oryza*, and *R^2^ =* 0.989, *p* =1.18 × 10^-12^, *N*= 211 for *Hordeum*; supplementary figure S5a). This significant correlation remained largely unchanged even after correcting deamination rates by subtracting baseline substitutions means (supplementary figure S5b). All samples that passed filtering showed the expected “mirrored” deamination patterns characteristic of aDNA (supplementary figure S5).

Both genera displayed statistically significant increases in the frequencies of 5’ C>T misincorporations at first position correlating with the age of the sample (figure 3d), with *Oryza* starting from a higher baseline of damage when compared to *Hordeum* and displaying a stronger relationship (*R^2^ =* 0.303, *p* = 8.62 × 10^-21^, *N*= 245 for *Oryza*, and *R^2^ =* 0.207, *p* =3.63 × 10^-12^, *N*= 211 for *Hordeum,* respectively). A slightly weaker yet significant relationship between 5’C>T substitutions and sample age was also observed when analysing all samples together (*R^2^ =* 0.106, *p* = 1.11 × 10^-12^, *N*= 456; supplementary figure S1d). In stark contrast to the significant relationship between 5’ C>T misincorporation and age, non-deamination substitution rates showed no significant relationship with sample age (supplementary figure S6), indicating that the patterns observed are specific to *post-mortem* deamination rates.

### Differences in rates of damage for *Hordeum* and *Oryza*

We compared differences in the regression slopes and intercepts for all aDNA damage metrics between the genera *Hordeum* and *Oryza* with an analysis of covariance (ANCOVA) and visualised this comparison with boxplots.

#### Endogenous content

The analysis of covariance revealed significant effects of both sample age (*Pr*(Sample age) = 0.0114, *N =* 456) and genus (*Pr*(Genus) = 0.0204, *N =* 456) on the fraction of endogenous DNA content (figure 5a). No significant interaction was observed between sample age and genus (*Pr*(Sample age: Genus) = 0.176, *N =* 456), indicating that the rate of exogenous DNA colonisation over time does not differ significantly between the two genera and is not time dependent. This was further supported by ANOVA between the model including the interaction and the model with no interaction, which confirmed that adding the interaction did not improve model fit (*F* = 1.841, *Pr =* 0.175). The model with no interaction displayed a significant fit (*F=* 5.395*, Pr =* 0.004835) but explained only a small proportion of the variance (*R^2^* = 0.02327), suggesting that factors beyond genus and sample age can substantially influence endogenous DNA content.

**Figure 5:**
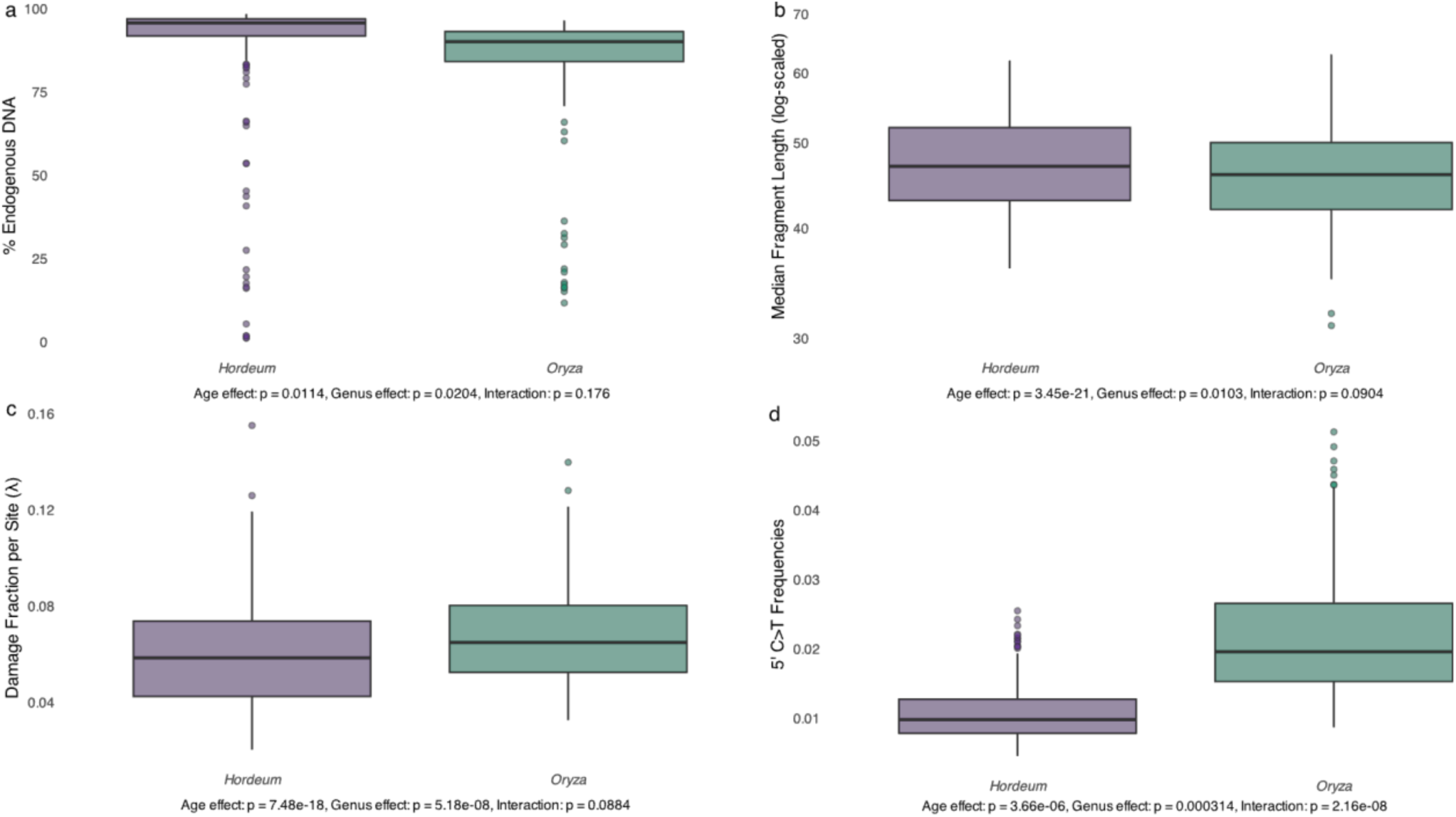
Analysis of Covariance (ANCOVA) of aDNA damage metrics for *Hordeum* and *Oryza*, with sample age as covariate and genus as factor. (a) Endogenous DNA content; (b) Fragment length (c) Damage fraction per site (λ); (d) 5’ C>T damage frequencies. Significance values reported below each boxplot indicate the effects of age, genus and their interaction upon the analysed aDNA damage metric.

#### Fragment length

The analysis of covariance revealed significant effects of sample age and genus on fragment length (figure 5b) (*Pr*(Sample age) < 3.45 × 10^-21^; *Pr*(Genus) = 0.0103; *N* = 456). The interaction between sample age and genus was not significant (*Pr*(Sample age: Genus) = 0.0893; *N* = 456) and its inclusion did not significantly affect model fit (*F* = 2.8993, *Pr* = 0.0904). The model with no interaction explained 18% of the variance in fragment length (*R^2^* = 0.1827) and displayed a significant fit (*F* = 50.65, *Pr* < 2.2 × 10^-16^), indicating that the genera differ in their baseline fragment lengths, but not in their rate of fragmentation over time.

#### Damage fraction per site (λ) and DNA decay rate (k)

We observed a slower DNA decay rate of *Oryza* (*k=* 1.79 × 10^-4^ per nucleotide per year) than that of *Hordeum* (*k=* 2.64 × 10^-4^ per nucleotide per year). The analysis of covariance revealed significant effects of both sample age (*Pr*(Sample age) = 7.48 × 10^-18^; *N* = 456) and genus (*Pr*(Genus) = 5.18 × 10^-8^; *N* = 456) on the rates of bond breaking (figure 5c). However, no interaction between sample age and genus was observed (*Pr*(Sample age: Genus) = 0.0884; *N* = 456) and including the interaction did not significantly improve model fit (*F* = 2.9152, *Pr* = 0.08843). The model with no interaction explained 18% of the variance in lambda values (*R²* = 0.1842) and displayed a highly significant fit (*F* = 51.14, *Pr* < 2.2 × 10⁻¹⁶). Therefore, the DNA decay rates (*k*) of *Hordeum* and *Oryza*, which correspond to the slopes of the regression between damage fraction per site (λ) and age, are not significantly different.

#### Nucleotide misincorporations

We observed a significant effect of both sample age (*Pr*(Sample age) = 3.66 × 10⁻⁶; *N* = 456) and genus (*Pr*(Genus) = 3.14 × 10⁻⁴; *N* = 456) on the rates of 5’ C>T misincorporations (figure 5d). The interaction between sample age and genus was significant (*Pr*(Sample age: Genus) = 2.16 × 10^-8^; *N* = 456), indicating that the rate of cytosine deamination over time differs substantially between the two genera. The model with interaction explained almost 55% of the variance in 5’C>T damage (*R²* = 0.5448) and displayed a significant fit (*F* = 180, *Pr* < 2.2 × 10⁻¹⁶). This was further corroborated by the ANOVA analysis, which confirmed that including the interaction significantly improved model fit (*F* = 32.49, *Pr* = 2.165 × 10⁻⁸). Therefore, not only do *Hordeum* and *Oryza* samples differ in their baseline levels of cytosine deamination, but they also accumulate this type of damage at significantly different rates over time, with *Oryza* showing a steeper increase in deamination with age.

### Effects of climatic variables upon rates of aDNA damage

We examined climatic effects using two complementary approaches: a “collection climate” model that assigned monthly temperature and precipitation values based on each sample’s recorded collection location and month, and an “annual climate” model that used annual means and seasonality measures based solely on geographical origin. Both models aimed at capturing climate-driven DNA damage occurring during the initial post-collection period (field handling, drying, and early preservation), rather than during subsequent long-term storage. This assumption is supported by two lines of evidence: first, the rapid initial phase of DNA degradation driven by endogenous nucleases and proteases and exogenous microbial digestion [51,52], which occurs immediately after host death, is known to dominate post-mortem DNA damage [2,6,7,25]. Second, herbarium specimens are subsequently stored under standardised conditions that minimise further environmental variation [2,25]. We further tested this by including herbarium identity (as a proxy for institutional storage conditions) as a factor, and indeed found negligible explanatory power for herbarium identity, supporting the premise that collection-time climate, rather than storage climate, is the relevant environmental predictor of DNA damage patterns. Variance partitioning analysis revealed distinct patterns of environmental and temporal control over different aspects of DNA preservation (figure 6). The relative importance of factors varied substantially among damage metrics, with some showing strong environmental sensitivity while others were primarily controlled by age-related processes. In addition to environmental factors, genus emerged as having significant explanatory power, possibly as a variable confounded with climate due to distinct distributions of the *Hordeum* and *Oryza* genera from temperate and tropical climates respectively (figure 2 and 6).

**Figure 6:**
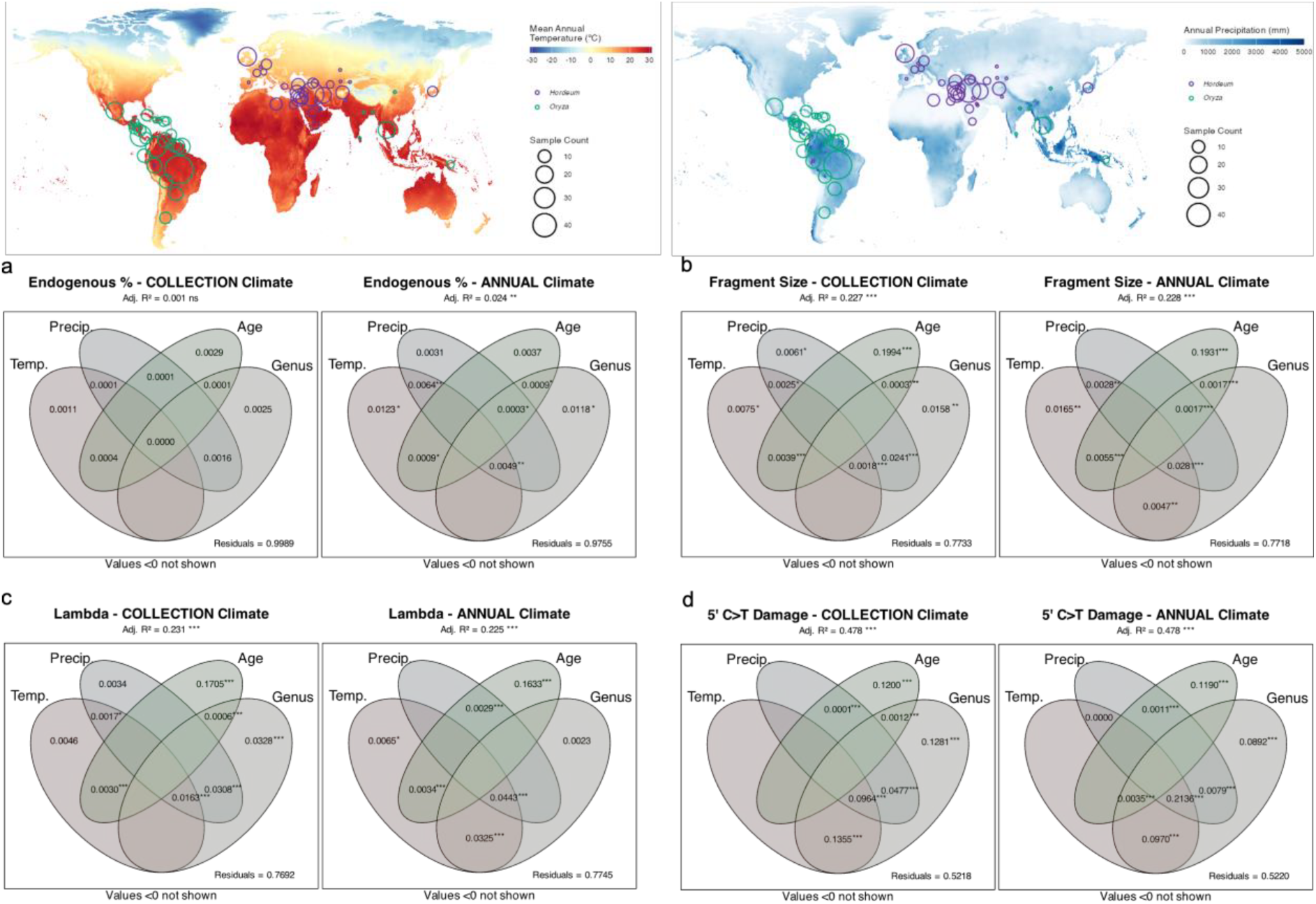
Climate influences on aDNA damage metrics in herbarium specimens. Maps (top panels) show the distribution of sampling locations for *Hordeum* and *Oryza* specimens overlaid on mean annual temperature (left) and annual precipitation (right) from the WorldClim climate dataset [53]. Venn diagrams (bottom panels) display the unique and shared contributions of the explanatory variables to the total variance in aDNA damage metrics.: (a) endogenous DNA fraction, (b) fragment size, (c) damage fraction per site (lambda), and (d) 5’ C>T substitution frequencies at first base. Each metric is analysed using two models: collection climate (left) and annual climate (right). Adjusted *R²* values for each model are shown above the plots. Asterisks indicate statistical significance for the overall models and for each unique predictor and combination of predictors (\**p* ≤ 0.05, \*\**p* ≤ 0.01, \*\*\**p* ≤ 0.001).

#### Endogenous content

Endogenous DNA content showed little association with the tested variables in the variance partitioning analysis (figure 6a). Only the collection climate model displayed statistical significance, but the fraction of variance explained by all analysed predictors was negligible (Adj. *R²* = 0.024, *p =* 0.007). The unique contributions of temperature and genus and the combined contributions of temperature, precipitation, age and genus showed marginal statistical significance (*p* ≤ 0.05, supplementary table S3a). However, their poor performance in explaining the fraction of variance (Adj. *R²* ranging from 0.001 to 0.01) indicates that endogenous DNA fraction is largely determined by factors not captured by climatic variable at the time of specimen collection nor age. Removing genus from the variance partitioning analysis did not significantly alter the results (supplementary figure S7a, supplementary table S4a). Similarly, inclusion of herbarium as a proxy of institutional preservation practices and post-collection long term storage conditions also did not significantly alter the results (supplementary figure S8a, supplementary table S5a).

#### Fragment length

Our models had more explanatory power when it came to fragment length, with both collection and annual climate factors being statistically significant (*p* ≤ 0.001) and performing similarly (figure 6b, Adj. *R²* = 0.227 and 0.228, respectively). Sample age was revealed as the largest contributing factor in both collection and annual models (Adj. *R²* = 0.199, 0.193 respectively), consistent with temporal degradation processes. The shared variance between climatic variables and age were small, indicating that age effects on fragment size are largely independent of environmental variables. Indeed, whilst climatic variables were significant in both models (supplementary table S3b), their unique and combined variance explained were negligible (Adj. *R²* ≤ 0.01). Removal of genus from the analysis marginally improved the unique and combined fraction of variance explained by temperature and precipitation in the collection and annual model respectively (supplementary figure S7b, supplementary table S4b), implying collinearity between genus and climatic variables, whereby further inclusion of herbarium had minimal effects (supplementary figure S8b, supplementary table S5b).

#### Damage fraction per site (λ)

Models predicting rates of DNA bond breaking (damage fraction per site, λ) from climatic and temporal variables explained up to 23% of the total variance in λ (figure 5c; Adj. *R²* = 0.231, 0.225 in the collection and annual models, respectively). Sample age contributed the largest unique fraction in both models (Adj. *R²* =: 0.170, 0.163), reflecting the fundamental relationship between specimen age and DNA backbone degradation. Temperature and precipitation showed minimal unique contributions and minimal shared variance with genus in the collection climate model (Adj. *R²* ranging from 0.01 to 0.03), and moderate shared variance with genus in the annual climate model (Adj. *R²* ranging from 0.03 to 0.04), suggesting indirect effects through correlations between genus and climatic variables. Indeed, removal of genus from the analysis did not strongly affect the overall variance explained by each model but inflated the unique and shared variance explained by temperature and precipitation (supplementary figure S7c).

#### Nucleotide misincorporations

The 5’ C>T damage metric showed the highest explained variance among all damage metrics, with both collection and annual climate models being significant (*p* ≤ 0.001) and explaining approximately 49% of the total variance (figure 6d). Whilst the unique contribution of age emerged as the strongest predictor of 5’ damage (Adj. *R²* = 0.120, 0.119 in the collection and annual models, respectively), most explanatory power derived from shared effects among predictors rather than unique contributions. The combined effects of temperature and genus explained a substantial proportion of the total variance (Adj. *R²* = 0.135, 0.097). A similar pattern was observed for the combined effect of precipitation and genus (Adj. *R²* = 0.047, 0.008) and for the combined effects of temperature, precipitation and genus (Adj. *R²* = 0.096, 0.213), suggesting a strong correlation between climatic variables and genus. Removal of genus from the analysis (supplementary figure S7d) revealed temperature as the largest unique contributing factor in the collection model (Adj. *R²* = 0.134), followed by sample age (Adj. *R²* = 0.121) and precipitation (Adj. *R²* = 0.047), whilst the combined effect of temperature and precipitation explained almost 10% of the variance (*R²* = 0.096). Notably, the variance explained by the combined effect of temperature and precipitation was inflated in the annual climate model (Adj. *R²* = 0.214, supplementary figure S7d) and emerged as the largest contributor, followed by unique contribution of sample age (Adj. *R²* = 0.109) and temperature (Adj. *R²* = 0.096). The inclusion of herbarium did not improve the total fraction of variance explained in neither model (supplementary figure S8d), with the unique and shared fraction of explained variance by herbarium and other variables (age, climate and genus) being negligible (Adj. *R²* ranging from 0.0003 to 0.01).

### Temperature effects on 5’ C>T deamination frequencies

Given the predictive power of temperature in explaining 5’ C>T damage patterns, we conducted post-hoc regression analyses to examine the relationships between temperature and cytosine deamination. We observed a strong positive relationship between temperature and 5’ C>T misincorporation frequencies (figure 7). The annual temperature model explained 18.4% of variance (*R^2^ =* 0.184, *p* = 1.93 × 10^-18^, *N*= 456, figure 7a), while the collection temperature model showed a slightly higher explanatory power (*R^2^ =* 0.201, *p* = 3.74 × 10^-20^, *N*= 456, figure 7b). However, this relationship was only observed when analysing the genera together and disappeared when analysing the genera on an individual basis (figure 7c, d).

**Figure 7:**
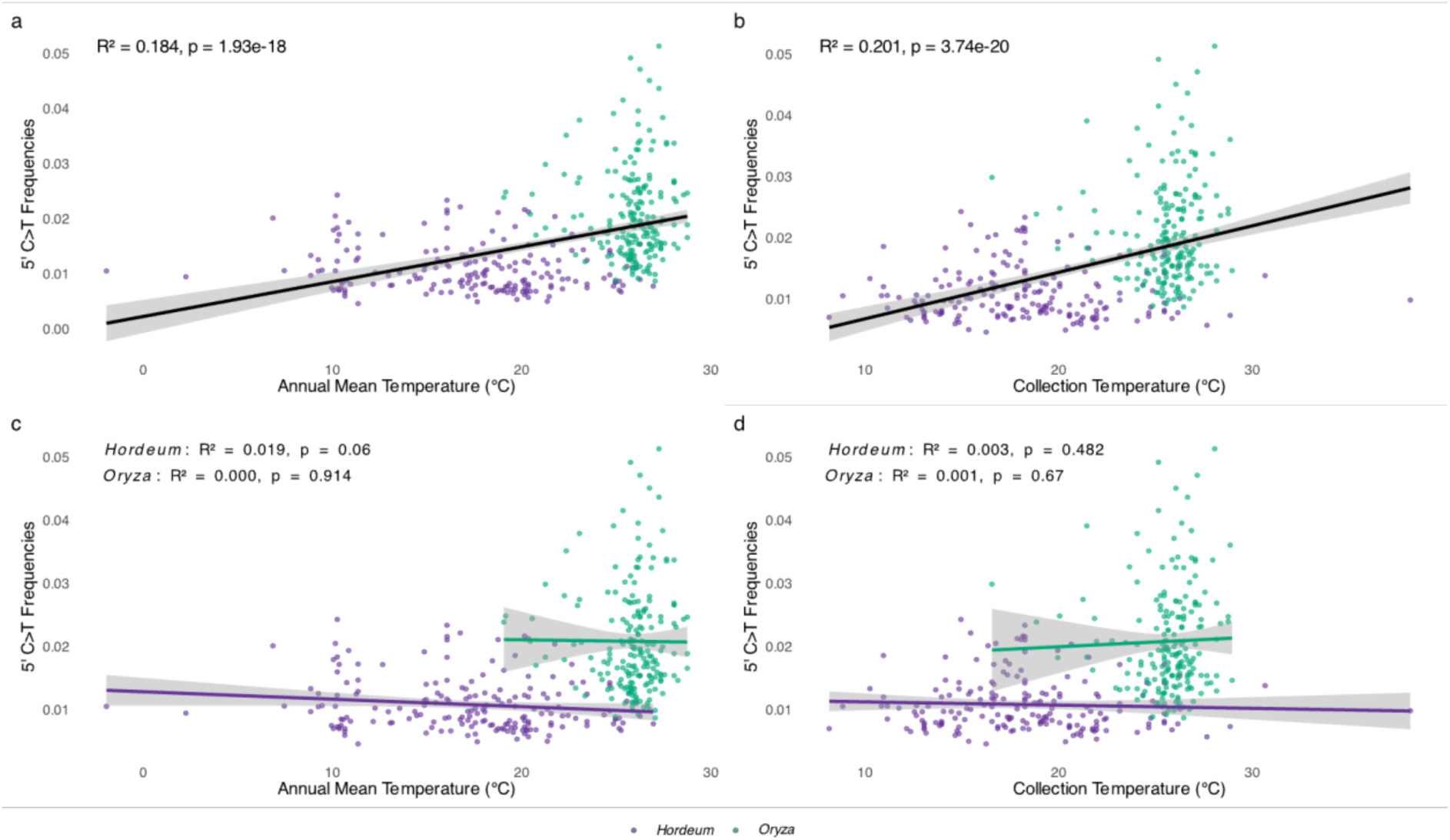
Relationship between temperature and deamination rates (5’ C>T misincorporation frequencies) in herbarium specimens for (a) annual mean temperature model, (b) collection temperature model, (c) annual mean temperature model for *Hordeum* and *Oryza*, and (d) collection temperature model for *Hordeum* and *Oryza*. Insets show regression statistics. The significant relationships observed when analysing all samples together (a, b) are largely driven by the contrasting climatic origins and damage levels between genera; these relationships are not significant when analysing each genus separately (c, d), indicating that temperature and genus effects are confounded in this dataset.

Furthermore, whilst the relationship between non-deamination substitutions and temperature showed some significance, the explanatory power was negligible (*R^2^ =* 0.007 - 0.07, supplementary figure S9), indicating that the temperature-damage relationship we reported are specific to deamination.

### Divergence-corrected nucleotide misincorporations analyses

All analyses of 5’ C>T misincorporations were repeated using deamination frequencies corrected for reads-reference divergence. After correction, mean 5’ C>T frequencies decreased from 1.74 ± 0.55% to 1.39 ± 0.50% across all samples (supplementary table S6). However, despite these corrections, all major patterns observed in the primary analyses remained largely unchanged. Corrected deamination frequencies still showed strong positive relationships with sample age in both genera (*R^2^ =* 0.318, *p* = 6.17 × 10^-22^, *N*= 245 for *Oryza*, and *R^2^ =* 0.205, *p* =4.61 × 10^-12^, *N*= 211 for *Hordeum,* respectively, supplementary figure S10), which were comparable to the uncorrected damage-age relationship observed in the primary analysis (*R^2^ =* 0.303, *p* = 8.62 × 10^-21^, *N*= 245 for *Oryza*, and *R^2^ =* 0.207, *p* =3.63 × 10^-12^, *N*= 211 for *Hordeum,* respectively, figure 3d). Variance partitioning analyses revealed age as the dominant factor explaining variation in corrected deamination rates (Adj. *R²* = 0.122, 0.156 in the collection and annual models, respectively), with the combined effects of temperature and genus contributing comparable fractions (Adj. *R²* = 0.110, 0.113, supplementary figure S11) to those obtained in the primary analyses (figure 6d). Similarly, the significant correlation between corrected deamination frequencies and temperature persisted for both annual (*R^2^ =* 0.171, *p* = 2.59 × 10^-18^, *N*= 456, supplementary figure S12a) and collection month models (*R^2^ =* 0.173, *p* = 2.74 × 10^-17^, *N*= 456, supplementary figure S12b) but disappeared when analysing the genera on an individual basis (supplementary figure S12c,d), which was also comparable to the results obtained in the primary analyses (figure 7).

## Discussion

Genomic inferences from preserved samples allows scientists to directly add temporal scales to the studies of evolutionary and ecological histories of species, and of fundamental questions about the mechanisms and tempo of evolution. The advent of high-throughput sequencing technologies fuelled the revolutionary growth of the field of archaeo- and paleo-genomics. In 2022, the number of ancient human genomes passed 10,000 with over 200 papers published [20]. The discovery of extremely well-preserved samples [54] and development of new approaches [55–58] have pushed the limits of DNA recovery and sequencing. For plants, owing to the great diversity of wild and cultivated taxa, it is not possible to know the exact number of sequenced preserved specimens. In recent publication of ‘Plant Tree of Life’ alone, genome-wide enrichment sequences have been generated for over 2,500 plant genera from herbarium specimens [59]. Current projects focusing on crops and their wild relatives routinely sequence whole genomes for hundreds of historical specimens [60–62]. RGB Kew is currently aiming to sequence genomes of 7,000 preserved specimens of fungi from its fungarium collection [63]. With increasing scale of aDNA research across all kingdoms, it is fundamentally important to improve our understanding of DNA preservation in historical and archaeological samples.

The focus of aDNA preservation research has been the impact of time on DNA degradation. Thanks to two well-researched examples, we indeed know that DNA deamination and depurination is correlated with sample age [13,25]. However, it has been noted that environmental conditions pre- and post-mortem should modulate the pace at which DNA is degraded [2]. Emphasis has been put on temperature and humidity, two parameters that have been suggested as reasons for difficulties in sequencing ancient genomes from the tropics [2]. Our work on herbarium specimens has allowed us to quantify the impact of age, environment and storage conditions on DNA preservation, revealing complex genus-specific differences extending beyond age-related degradation. Our regression analyses showed relationships between sample age and different damage metrics, suggesting that while some aspects of aDNA degradation follow predictable temporal trajectories, others are more heavily influenced by environmental and biological factors.

### Age-dependent aDNA degradation patterns

We further confirmed the highly fragmented nature of aDNA retrieved from herbarium specimens, with the median fragment size of all analysed samples averaging 46.24 bp (SD = 6.97). This is consistent with previous studies on herbarium samples of a similar age range to that of the current study [25] and on dry-pinned arthropods museum collections [9], but also comparable to fragment sizes observed for animal remains that are several orders of magnitude older, from a few hundred up to thousands of years old [5,13]. Possibly owing to lower levels of environmental variation experienced by herbarium samples [25], we were able to detect a weak yet significant relationship between median fragment length and collection year (figure 3b). For a subset of samples (*N* = 40), we directly measured gDNA fragment size distributions prior to library preparation with high-resolution capillary electrophoresis (TapeStation), allowing us to assess DNA fragmentation independent of any library construction artifacts or bias. When examining the relationship between gDNA fragment size (TapeStation) and collection year, we found an even stronger correlation with age (figure 4). This contrasts with findings from animal bone studies, where no correlation between DNA fragmentation and sample age has been observed [5,13,64], even when controlling for environmental variables [2]. This fundamental difference can be explained by the two-step process of DNA decay and degradation: a first rapid phase of enzymatic decomposition occurring immediately after host death, driven by endogenous nucleases and proteases [51], and microbial digestion [52]; and a second phase of chemical decomposition driven by hydrolytic and oxidative reactions, occurring at much lower rates [6,7]. While the environment during sampling of herbarium specimens is variable, the standardized preparation and storage procedures used in herbaria reduce environmental variation during storage compared to the highly variable burial conditions experienced by animal bones. Indeed, the source herbaria (a proxy for storage conditions) have very limited explanatory power for DNA degradation (figure 6). The lower levels of environmental variation experienced by herbarium specimens allows detection of the underlying temporal fragmentation process that occurs during the second phase of chemical degradation [25,51]. In archaeological contexts, the influence of variable environmental, physical and chemical conditions, as well as tissue types and sample excavation and storage [2,13], may mask more subtle age-related fragmentation patterns that become detectable under the controlled conditions of herbarium storage [25].

The distinct preservation pathways of herbarium specimens compared to bones and museum specimens also likely reflect different balances between hydrolytic and oxidative DNA degradation processes. Herbarium specimens are often dried with heat upon collection [43,65]. Whilst this rapid desiccation can curtail hydrolytic damage, it also increases rates of oxidative processes [6,43,65]. In contrast, animal bones or museum specimens such as pinned arthropods dry more slowly, either through gradual burial diagenesis or museum preparation. These mechanistic differences can contribute to the variation in DNA decay rates observed across different biological materials: our herbarium samples exhibited decay rates (*k* = 2.08 × 10⁻⁴) nearly eight times faster than that observed in bones (*k* = 2.71 × 10⁻⁵;[13]), suggesting a higher susceptibility of *post-mortem* enzymatic and chemical DNA damage in herbarium samples compared to bones [25]. Interestingly, arthropod museum specimens show even faster decay (*k* = 4.6 × 10⁻⁴,[9]), approximately twice the rate observed in our herbarium samples. This variation likely reflects differences in tissue composition (cellulose/lignin in plants, chitin/protein in arthropods and hydroxyapatite/collagen in bones), storage methods (pressed herbarium sheets, pinned insects, and buried remains), and the distinct degradation dynamics imposed by rapid versus gradual desiccation. The intermediate position of herbarium specimens between museum arthropods collections and ancient bones in terms of decay rate suggests that DNA fragmentation is influenced by both intrinsic tissue properties and storage conditions.

Furthermore, the 5’ C>T damage patterns also showed significant correlations with age in both genera (figure 3d). However, while both genera show similar rates of DNA fragmentation, their divergent responses to chemical modifications suggest different susceptibilities to oxidative damage and deamination. The higher baseline damage and steeper accumulation rates in *Oryza* species indicate that the genus is more susceptible to *post-mortem* deamination, possibly due them growing in different environments.

While aDNA damage metrics such as fragmentation and deamination were observed to be time-dependent processes, the fraction of endogenous DNA content showed no correlation with age (figure 3a), suggesting that microbial colonization and subsequent displacement of endogenous DNA is largely independent of specimen age. Biological degradation processes are thus primarily driven by factors other than time, such as *ante-* and *post-mortem* microbial colonisation, tissue characteristics, and individual handling and storage conditions [66]. However, whilst we used species-specific reference genomes, increasing evolutionary distance between query and reference might have impacted this metric [9]. We therefore suggest caution when interpreting such results.

### Environmental controls on aDNA preservation

Our variance partitioning analysis shows that climatic effects on aDNA damage are largely mediated through complex interactions among predictors rather than independent contributions. The dominance of shared variance fractions indicates that temperature, precipitation and genus effects are highly correlated in our dataset, reflecting the geographic and temporal sampling patterns of herbarium collections.

For metrics reflecting DNA backbone integrity (fragment size and lambda), age consistently contributed the largest unique variance fractions, confirming that temporal degradation processes are the primary drivers of DNA fragmentation. The relatively large unique age contributions suggest that these physical degradation processes proceed independently of environmental and taxonomic factors once specimens enter standardized herbarium preparation methods storage practices. This age-dependent fragmentation appears to be a distinctive feature of museum specimens (such as herbarium specimens [25] and dry-pinned arthropods collections [9]) that contrasts with the patterns observed in archaeological animal bones [5,13]. For the latter, variable burial or storage conditions introduce environmental heterogeneity that can mask the underlying temporal degradation signal, whereas the standardised preparation and storage practices in museum specimens reduce such variation, allowing age-related fragmentation patterns to emerge.

The strong correlation between temperature and 5’ C>T damage provides mechanistic insight into why *Oryza* specimens, predominantly from tropical and sub-tropical regions, consistently show higher damage levels compared to temperate *Hordeum* specimens (figure 7). Elevated temperatures accelerate the spontaneous deamination of cytosine residues, explaining both the higher baseline damage observed in *Oryza* and the overall temperature-damage relationship observed across all herbarium samples. Precipitation variables as a proxy of humidity also showed significant effects, indicating that deamination is a time-dependent process modulated by temperature and humidity [2]. Overall, the complex interplay between temperature, humidity and their seasonality and damage metrics suggests that specimens from regions with less seasonal climates may experience different preservation trajectories than those from highly seasonal climates. The substantial unique genus contributions to 5’ C>T damage confirm differential susceptibility to cytosine deamination between *Hordeum* and *Oryza*. However, the large shared variance fractions between genus and temperature reflect confounding effect of the contrasting geographic origins of these genera (temperate vs. tropical; figure 6). This geographic segregation limits our ability to fully disentangle genus-specific susceptibility from temperature-driven effects upon DNA damage. When examining temperature-damage relationships within each genus separately (figure 7c, d), the correlations become non-significant. It is possible that genus captures other environmental factors that were not included in temperature and precipitation. Precipitation is a decent proxy for humidity, but there are multiple other factors influencing it, and high humidity is expected to accelerate the deamination process. Alternatively, the differences observed due to tropical and temperate samples (captured in genus) could be explained by different sample processing approaches in the two areas, with tropical specimens being more often oven-dried/baked. Indeed, baking has been shown to substantially affect DNA degradation [65]. Additionally, in the tropics, alcohol treatment used to be common to prevent moulding, and this has been shown to limit the success in DNA amplification, presumably due to rapid degradation [67]. Future studies incorporating additional plant families with overlapping geographic distributions would be necessary to clearly separate genus-specific susceptibility from environmental temperature effects. Nevertheless, the consistent genus-level difference we observe remains informative about the challenges of DNA preservation across different biogeographic contexts.

The negligible variance explained for endogenous fraction across all models supports our hypothesis that microbial colonization is largely independent of the measured environmental and biological predictors. This finding suggests that endogenous DNA loss is driven by stochastic factors such as initial contamination loads, handling procedures, or unmeasured specimen-specific characteristics.

### Conclusions and implications for herbarium curation and ancient DNA research

We show that ancient DNA damage patterns result from complex interactions between temporal degradation processes, environmental conditions during sampling, biological properties and storage condition. Herbarium specimens exhibit age-dependent DNA fragmentation patterns that are not observed in animal bone studies, indicating that standardized preservation conditions can reveal underlying temporal degradation processes masked by environmental variation in archaeological contexts. We identified significant genus-specific differences in aDNA damage susceptibility and established temperature as the dominant environmental driver of cytosine deamination.

Environmental effects on DNA damage operate primarily through complex interactions with taxonomic and temporal factors rather than direct independent contributions. Age-related degradation dominates physical DNA breakdown, while genus-specific differences in cytosine deamination susceptibility reflect both environmental adaptation and intrinsic biochemical properties. The comparison across biological materials reveals that tissue composition fundamentally constrains preservation potential, while storage conditions modulate decay rates.

These findings highlight the importance of considering specimen age, climatic origin and storage conditions when selecting herbarium specimens for ancient DNA analyses. The mechanisms underlying genus-specific differences in aDNA damage accumulation warrant detailed biochemical investigation. The significant, although small effect of herbarium-specific factors on damage metrics indicates that institutional preservation practices vary and represent an underexplored factor in ancient DNA research. Future studies should provide insights into the biological processes driving DNA degradation, including detailed microbiome analyses. Understanding which microbial taxa are most problematic for DNA preservation and how their colonization is influenced by specimen characteristics could inform targeted preservation strategies. As the scale of ancient DNA research continues to expand, such findings will be essential for maximizing the scientific value of the world’s natural collections and informing evidence-based approaches to specimen sampling and preservation.

## Availability of code and requirements

Project name: Herbaria aDNA Damage

Project home page: https://github.com/Stefano-Porrelli/Herbaria_aDNA_Damage

Operating system(s): Linux/Unix (tested on SLURM-based HPC systems)

Programming language: Bash, R (version ≥4.0)

Other requirements:

- Conda/Miniconda,
- AdapterRemoval (≥2.0), BWA (≥0.7.17), FastQC (≥0.11), SAMtools (≥1.10), DeDup (≥0.12), mapDamage2 (≥2.0), MultiQC (≥1.8), Preseq (≥2.0), AMBER.
- R packages: dplyr, tidyr, purrr, stringr, readr, MASS, vegan, car, ggplot2, colorspace, viridis, ggrepel, ggtext, ggpubr, gridExtra, cowplot, sf, geodata, terra, maps
- External data: CHELSA V2.1 climate data (https://chelsa-climate.org/)

License: MIT licence

Any restrictions to use by non-academics: None

## Data availability

Raw FASTQ DNA sequences of all samples processed at the Royal Botanic Gardens, Kew (UK) are deposited on the sequence reads archive (SRA) on NCBI: *Hordeum vulgare* sequences are deposited under BioProject PRJNA1288534. *Hordeum spontaneum* sequences are deposited under BioProject PRJNA1289164. *Oryza rufipogon* sequences are deposited under BioProject PRJNA1288425. *Oryza grandiglumis* sequences are deposited under BioProject PRJNA1288424. *Oryza latifolia* sequences are deposited under BioProject PRJNA1288423. All remaining raw FASTQ DNA sequences of *Oryza* are deposited on the SRA under BioProject PRJNA1302186. Scripts and dataset to reproduce the analyses are available at https://github.com/Stefano-Porrelli/Herbaria_aDNA_Damage.

## Authors’ contributions

S.P. and R.M.G. conceived the project and designed the study; R.M.G. and S.P. designed the sampling strategy, with contributions from A.F. and P.H.L.; P.J.K and N.S. overseen and supervised the generation of sequencing data for *Hordeum* samples; S.P. sampled herbaria and performed the aDNA laboratory work and bioinformatic screening for the *Hordeum* samples processed at RBGK; A.H. sequenced the *Hordeum* libraries at IPK; P.H.L. sampled herbarium samples and performed the aDNA laboratory work and bioinformatic screening for the *Oryza* samples processed at RBGK; A.F. selected herbarium specimens, A.F., M.N.R., and R.A.W. performed the sampling, A.F., W.Y., N.M., M.N.R., and A.C.C. isolated aDNA and prepared aDNA libraries, and A.F. performed screening and validation for the *Oryza* samples processed at KAUST/UoN; S.P. analysed the historical data and interpreted the results with the contribution of R.M.G. and P.H.L; S.P. and R.M.G. wrote the manuscript with contributions from all authors.

## Acknowledgements

We thank curators for providing herbarium specimens: Anna Haigh and Sue Zmartzy (Royal Botanic Gardens, Kew); Paul Peterson and Robert Soreng (Smithsonian Institute Herbarium); Jordan K. Teisher (Missouri Botanic Gardens); Miriam Gaudeul (National History Museum, France); Michaela Schmull and Anthony R. Brach (Harvard Herbarium); Matthew C. Pace (New York Botanic Gardens). We acknowledge the expert technical assistance of Ines Walde and Jacqueline Pohl during DNA sequencing of *Hordeum* samples at the Leibniz Institute of Plant Genetics and Crop Plant Research (IPK). We also thank Anne Fiebig for her expert technical assistance with data submission of *Hordeum* low-throughput SRA sequences at IPK.

## Ethics and permissions

Historical herbarium specimens were obtained from established herbaria following institutional specimen access protocols. All sampling was conducted under standard museum loan agreements and followed institutional guidelines for destructive sampling of herbarium material. No additional ethical permissions were required as the study involved only preserved specimens collected under historical botanical collecting practices prior to CBD, but authors adhere to the ethical principles for Access and Benefit Sharing.

