## Supplementary Figures for "Patterns of aDNA Damage Through Time and Environments – lessons from herbarium specimens"

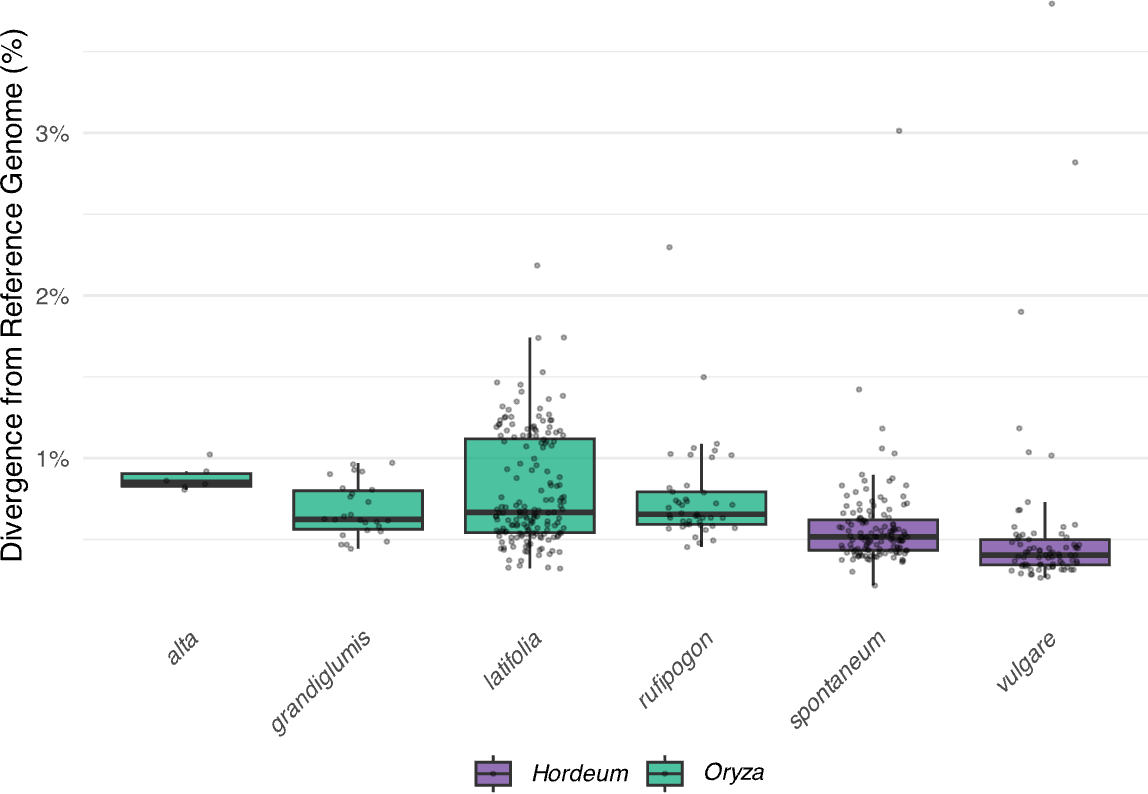

Supplementary figure S1: Evolutionary divergence between each species included in this study and their relative reference genome used for mapping. Divergence rates were inferred from baseline substitution rates (calculated as the mean frequency of non-deamination substitution types at position 1 from both 5' and 3' ends).

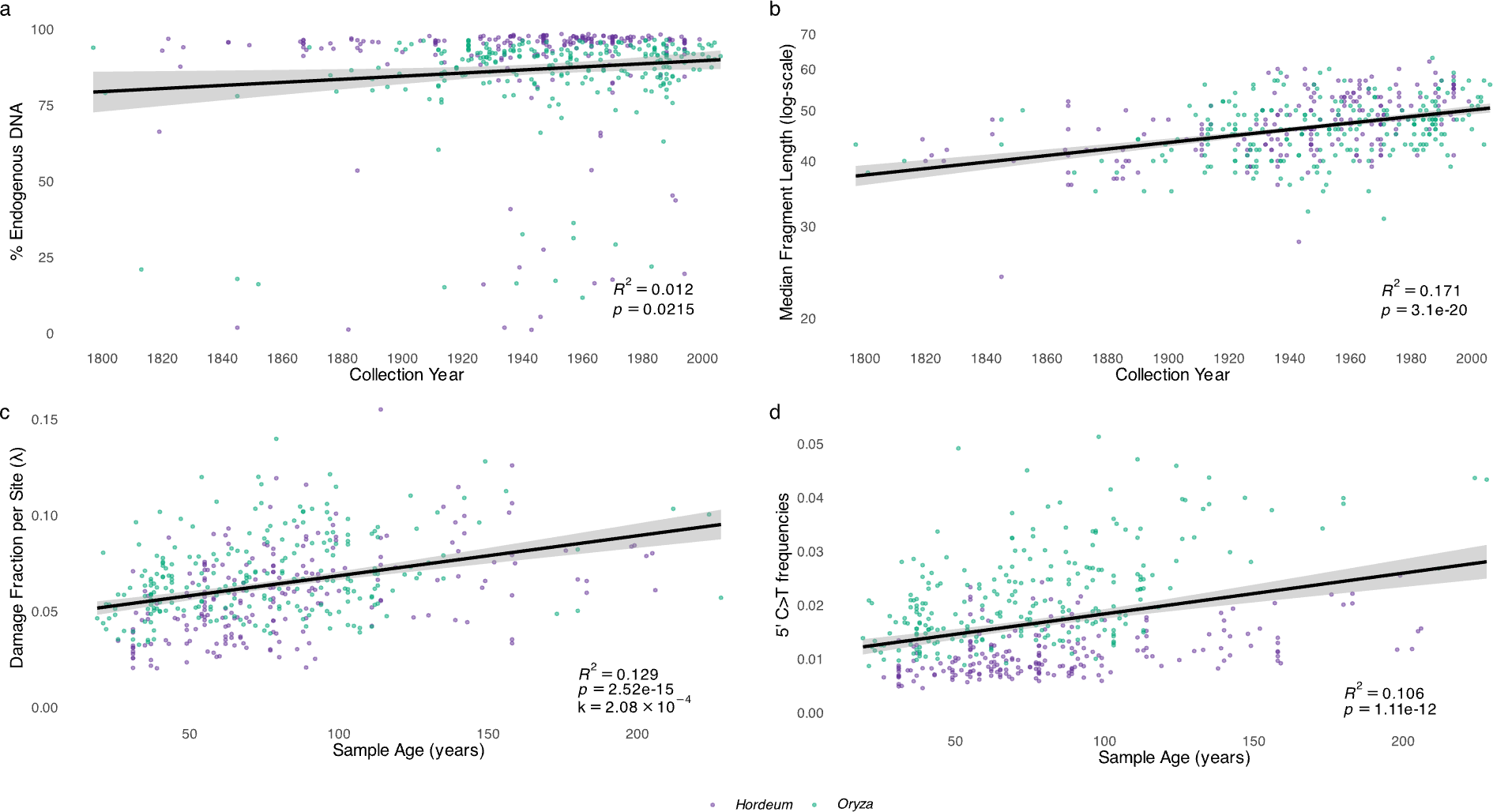

Supplementary figure S2: Regression analyses of aDNA damage metrics for all herbarium samples: (a) Fraction of endogenous DNA as a function of collection year. (b) Median fragment length of merged reads as a function of collection year, with log-scaled y-axis to show exponential relationship. (c) Damage fraction per site (λ) as a function of sample age, with the slope of regression corresponding to the DNA decay rate per base per year for all samples (*k* = 2.08 x 10^-4^). (d) Frequencies of C>T misincorporations at first base (5’ -end) as a function of sample age. Insets show regression statistics for each aDNA damage metric.

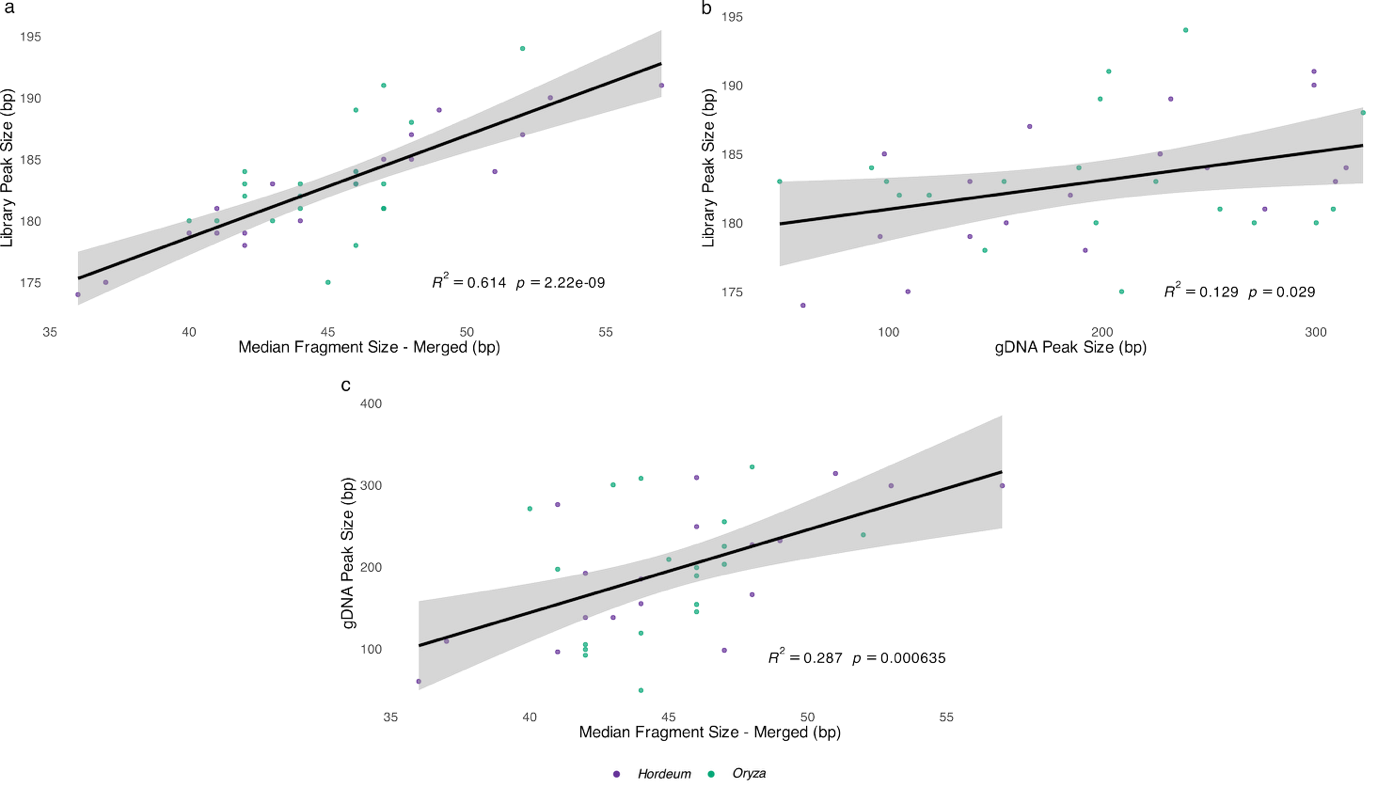
Supplementary figure S3: Regression analyses for a subset of herbarium samples (*N* = 40) between: (a) Peak size of libraries and median fragment size of the merged reads. (b) Peak size of libraries and genomic DNA. (c) Peak size of genomic DNA and median fragment size of merged reads. Insets show regression statistics.

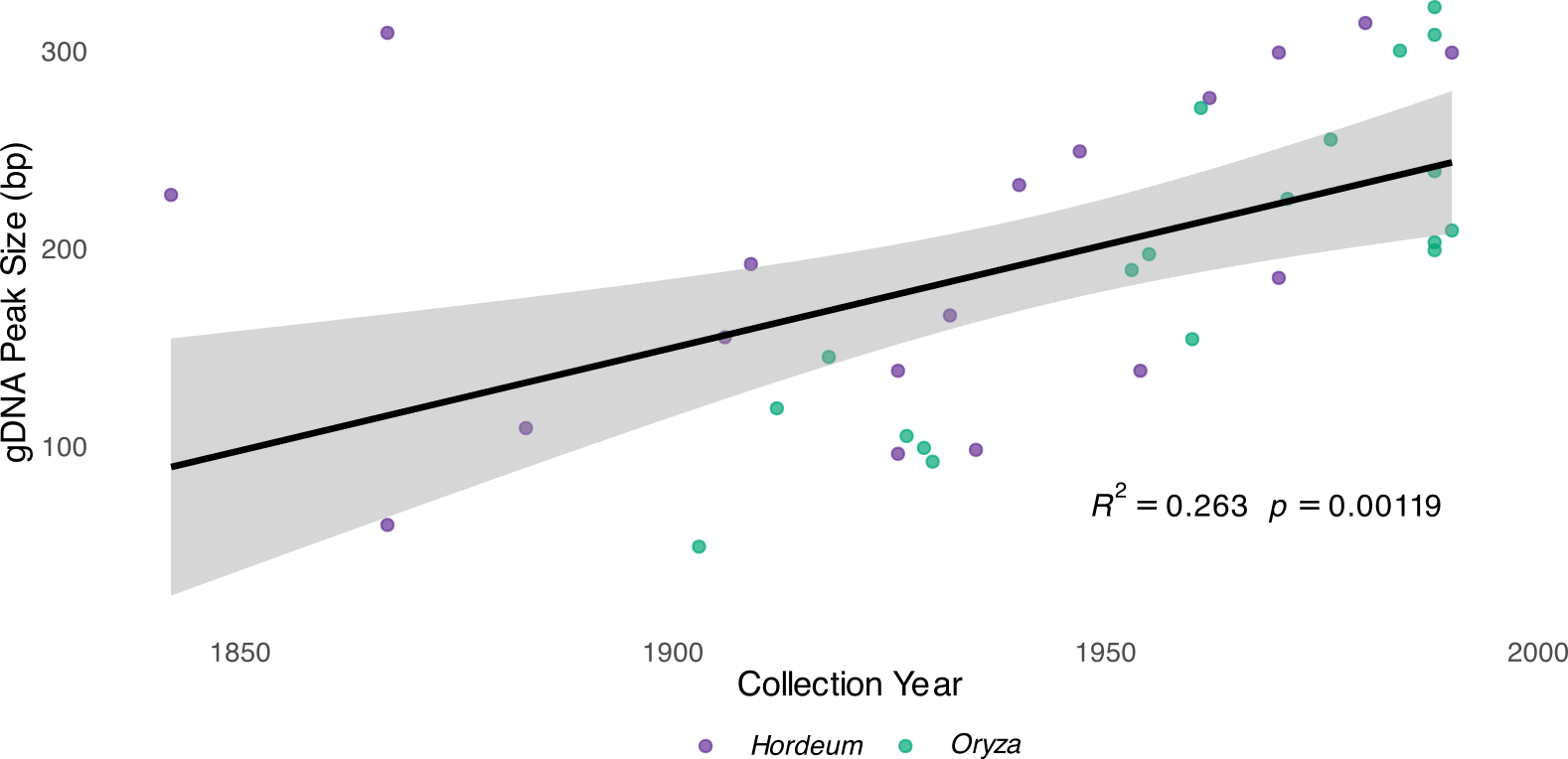

Supplementary figure S4: Regression between peaks of genomic DNA (gDNA) fragment size distribution and collection year for a subset of samples (*N* = 40) with inclusion of outliers.

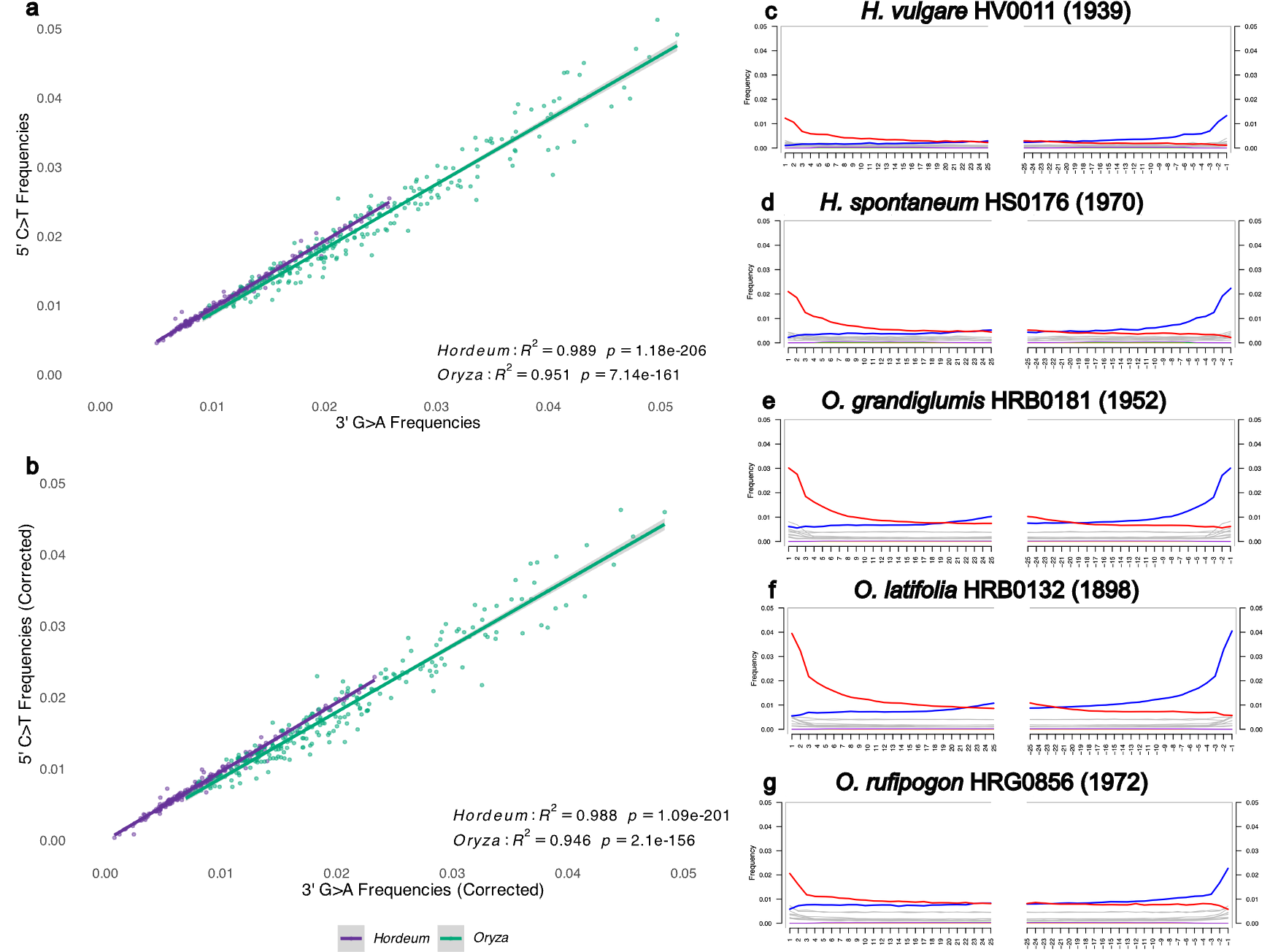

Supplementary figure S5: Left panels show (a) regression between C>T misincorporation frequencies at 5’ first base and G>A misincorporation frequencies at 3’ first base in double-stranded libraires, and (b) regression between C>T misincorporation frequencies at 5’ first base and G>A misincorporation frequencies at 3’ first base in double-stranded libraires after correcting deamination rates by subtracting mean baseline substitutions. Both uncorrected and corrected deamination rates show a significant and strong correlation for both genera. Right panels show examples of MapDamage2 misincorporation spectra displaying exponential increments of 5’ C>T frequencies (left plot, red line) and the complementary exponential increments of 3’ G>A (right plot, blue line) diagnostic of aDNA authenticity for (c) *H. vulgare* sample HV0011 collected in 1939, (d) *H. spontaneum* sample HS0176 collected in 1970, (e) *O. grandiglumis* sample HRB0181 collected in 1952, (f) *O. latifolia* sample HRB0132 collected in 1898, and (g) *O. rufipogon* sample HRG0856 collected in 1972.

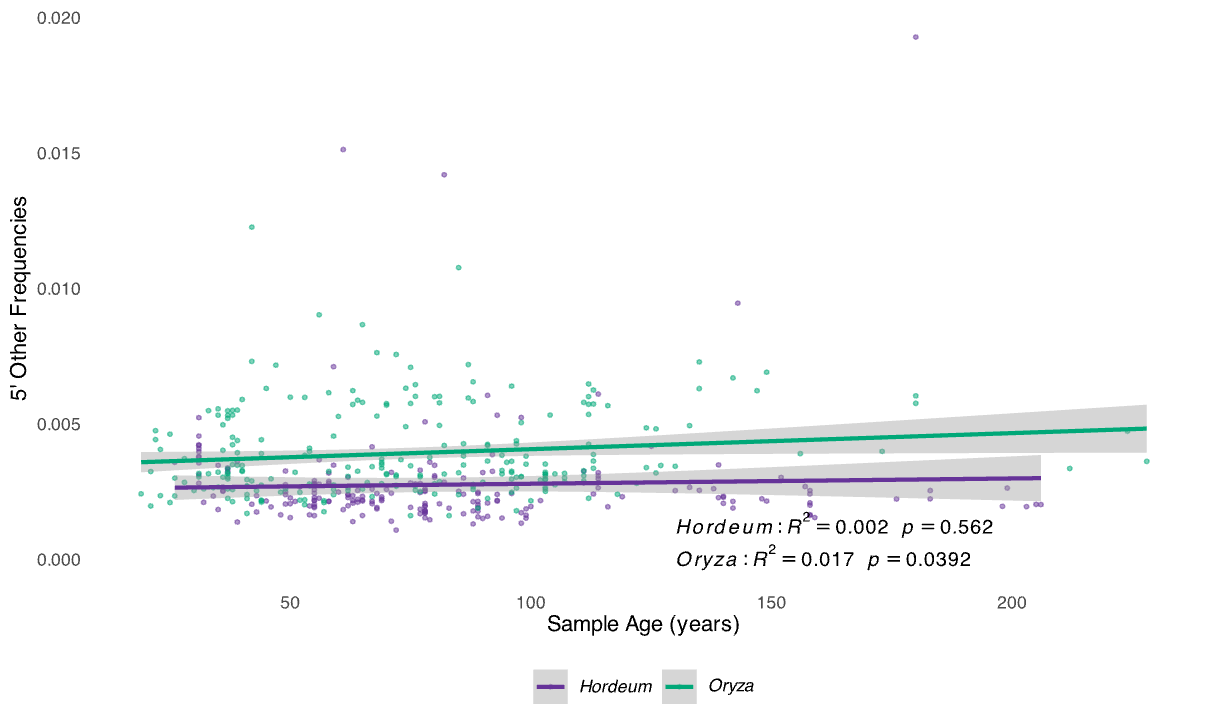

Supplementary S6: Relationship of non-deamination substitution frequencies at 5’ and sample age for *Hordeum* and *Oryza*. The significant relationship between deamination rates and age we observed is not reflected for non-deamination substitution rates, indicating that the patterns reported are specific to deamination frequencies, rather than mapping artefacts or read-reference divergence.

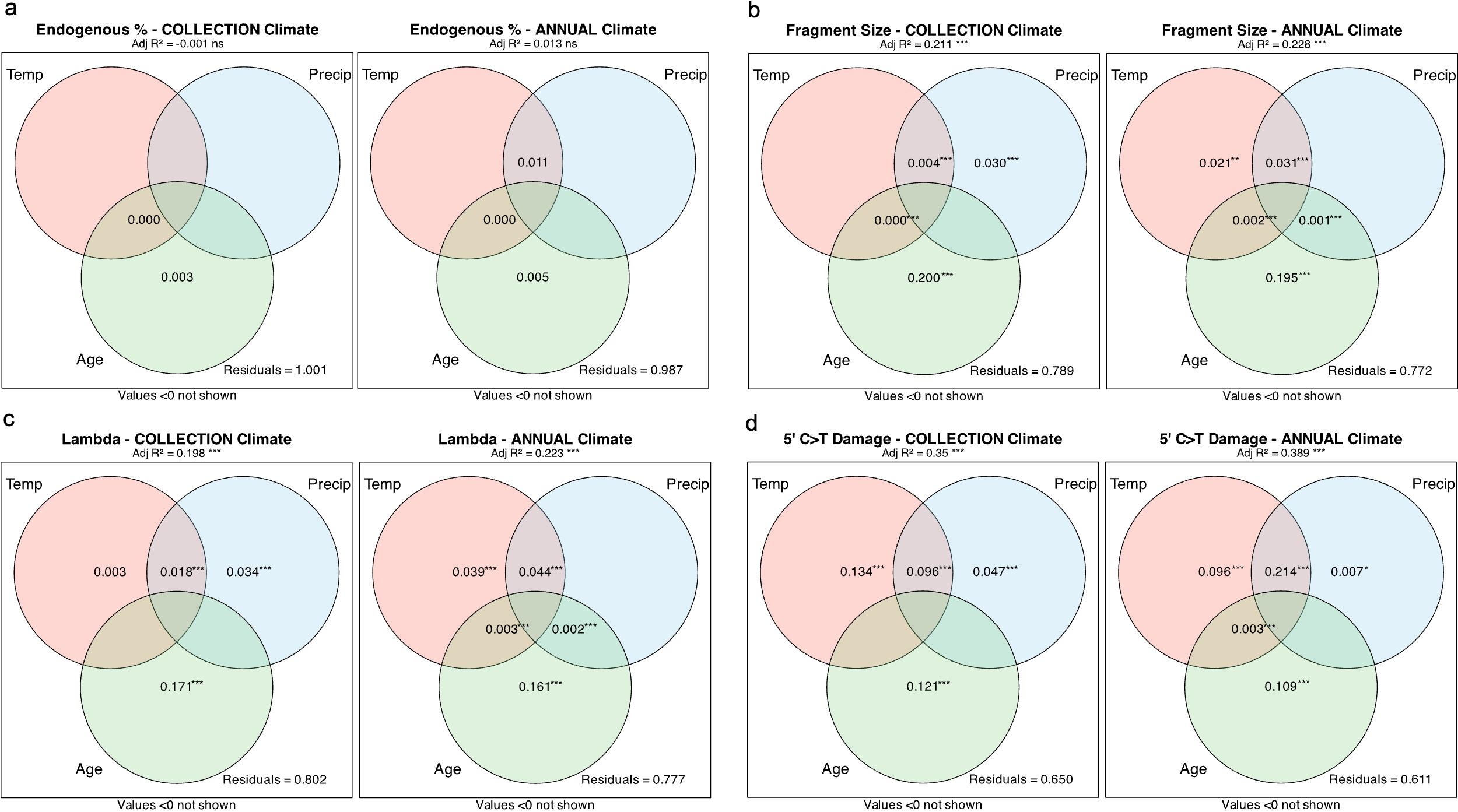

Supplementary figure S7: Climate influences on aDNA damage metrics in herbarium specimens. Venn diagrams display the unique and shared contributions of the explanatory variables (temperature, precipitation and age) to the total variance in aDNA damage metrics.: (a) endogenous DNA fraction, (b) fragment size, (c) damage fraction per site (lambda), and (d) 5' C>T substitution frequencies at first base. Each metric is analysed using two models: collection climate (left) and annual climate (right). Adjusted *R²* values for each model are shown above the plots. Asterisks indicate statistical significance for the overall models and for each unique predictor and combination of predictors (**p* ≤ 0.05, ***p* ≤ 0.01, ****p* ≤ 0.001).

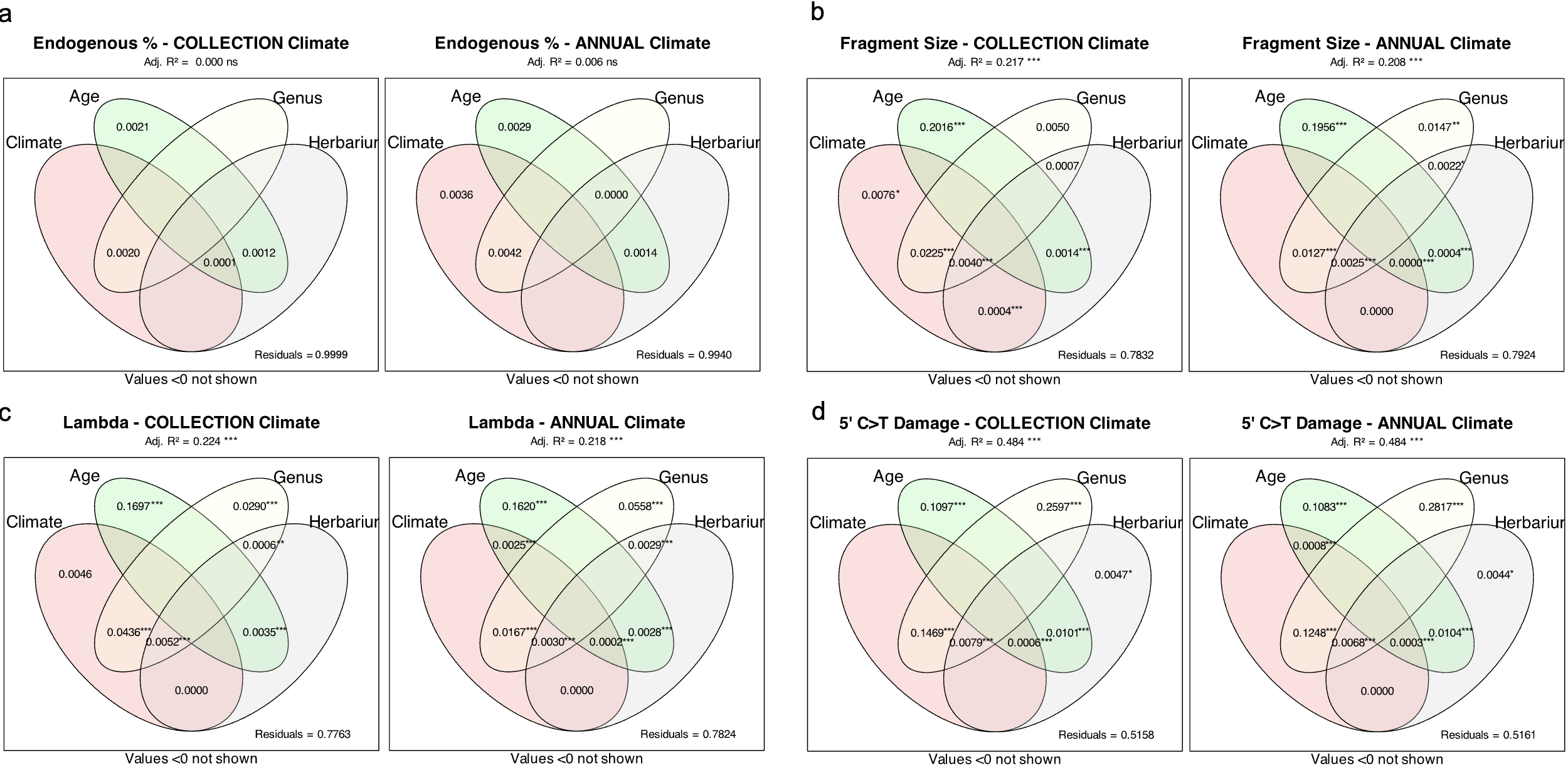

Supplementary figure S8: Climate influences on aDNA damage metrics in herbarium specimens. Venn diagrams display the unique and shared contributions of the explanatory variables (climate, age, genus and herbarium) to the total variance in aDNA damage metrics.: (a) endogenous DNA fraction, (b) fragment size, (c) damage fraction per site (lambda), and (d) 5' C>T substitution frequencies at first base. Each metric is analysed using two models: collection climate (left) and annual climate (right). Adjusted *R²* values for each model are shown above the plots. Asterisks indicate statistical significance for the overall models and for each unique predictor and combination of predictors (**p* ≤ 0.05, ***p* ≤ 0.01, ****p* ≤ 0.001).

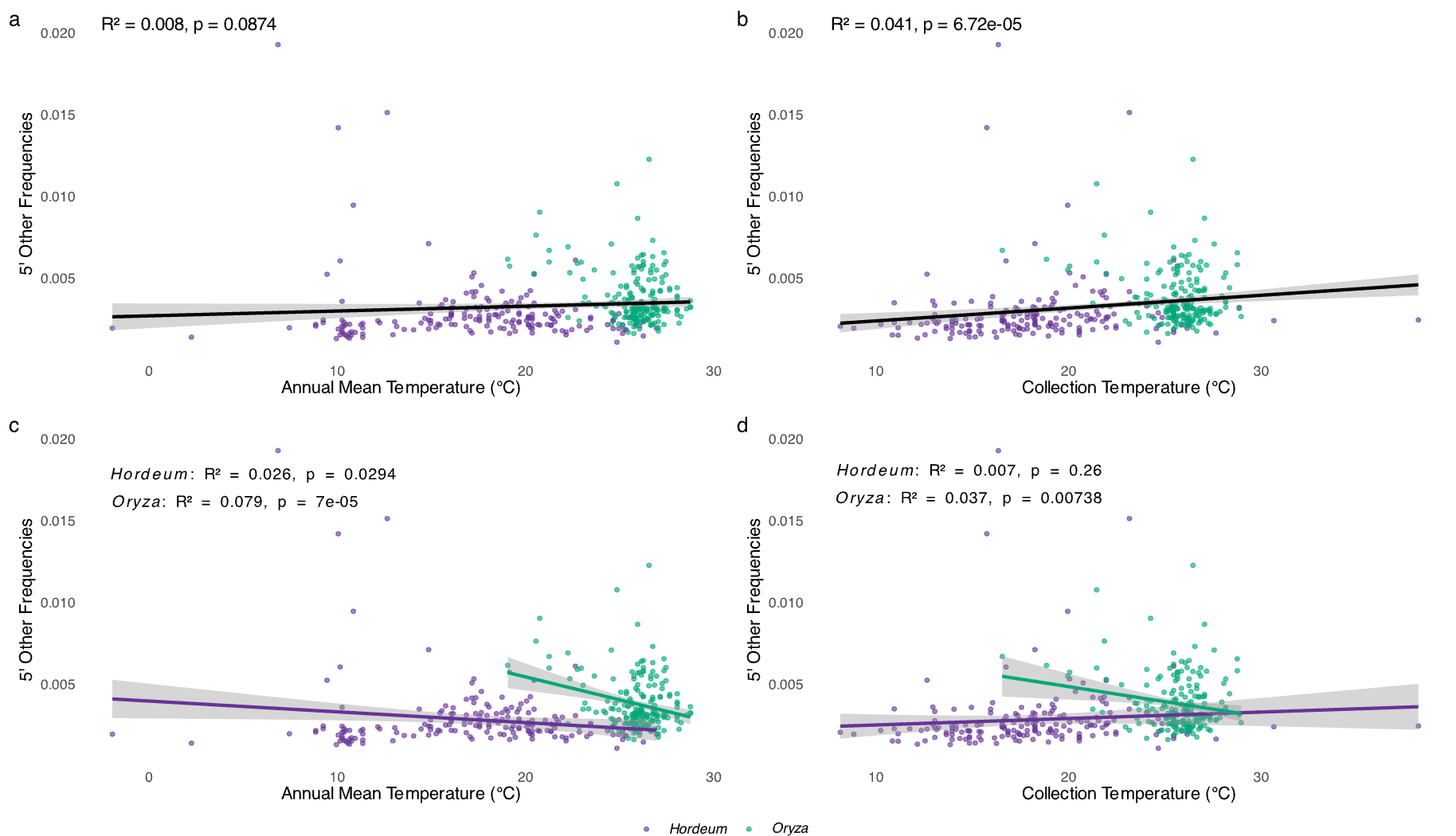

Supplementary figure S9: Relationship between temperature and baseline substitution rates in herbarium specimens for (a) annual mean temperature model, (b) collection temperature model, (c) annual mean temperature model for *Hordeum* and *Oryza*, and (d) collection temperature model for *Hordeum* and *Oryza*. Insets show regression statistics. Whilst the relationship between non-deamination substitutions and temperature showed some statistical significance (panel b and panels c,d for *Oryza* samples), the explanatory power was negligible, indicating that the temperature-damage relationship reported in the primary analyses are specific to deamination and are largely unaffected by read-reference divergence.

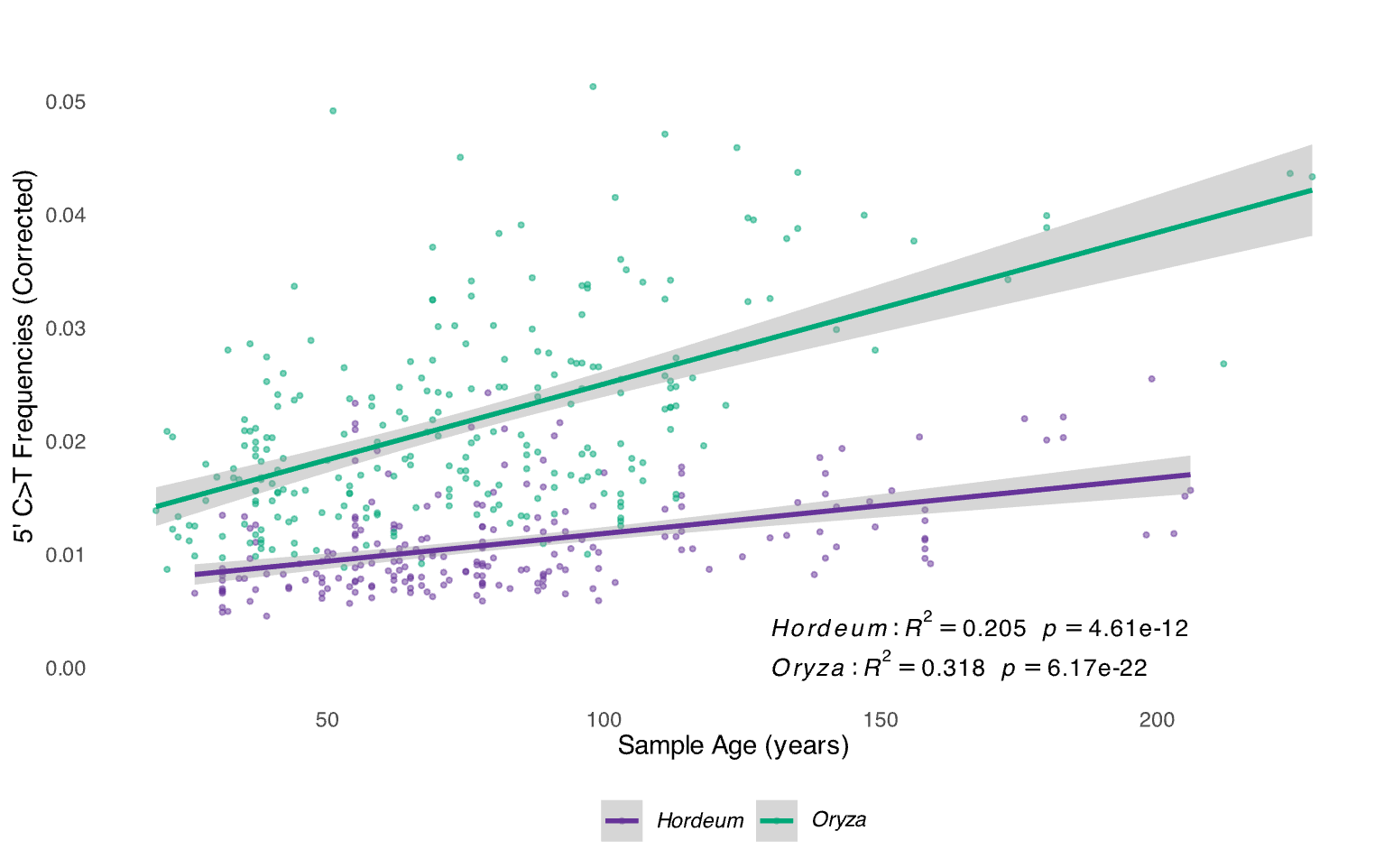

Supplementary figure S10: Regression analysis of C>T misincorporations frequencies at first base (5’ -end) for *Hordeum* and *Oryza* (corrected by baseline substitutions) as a function of sample age. Insets show regression statistics for each genus.

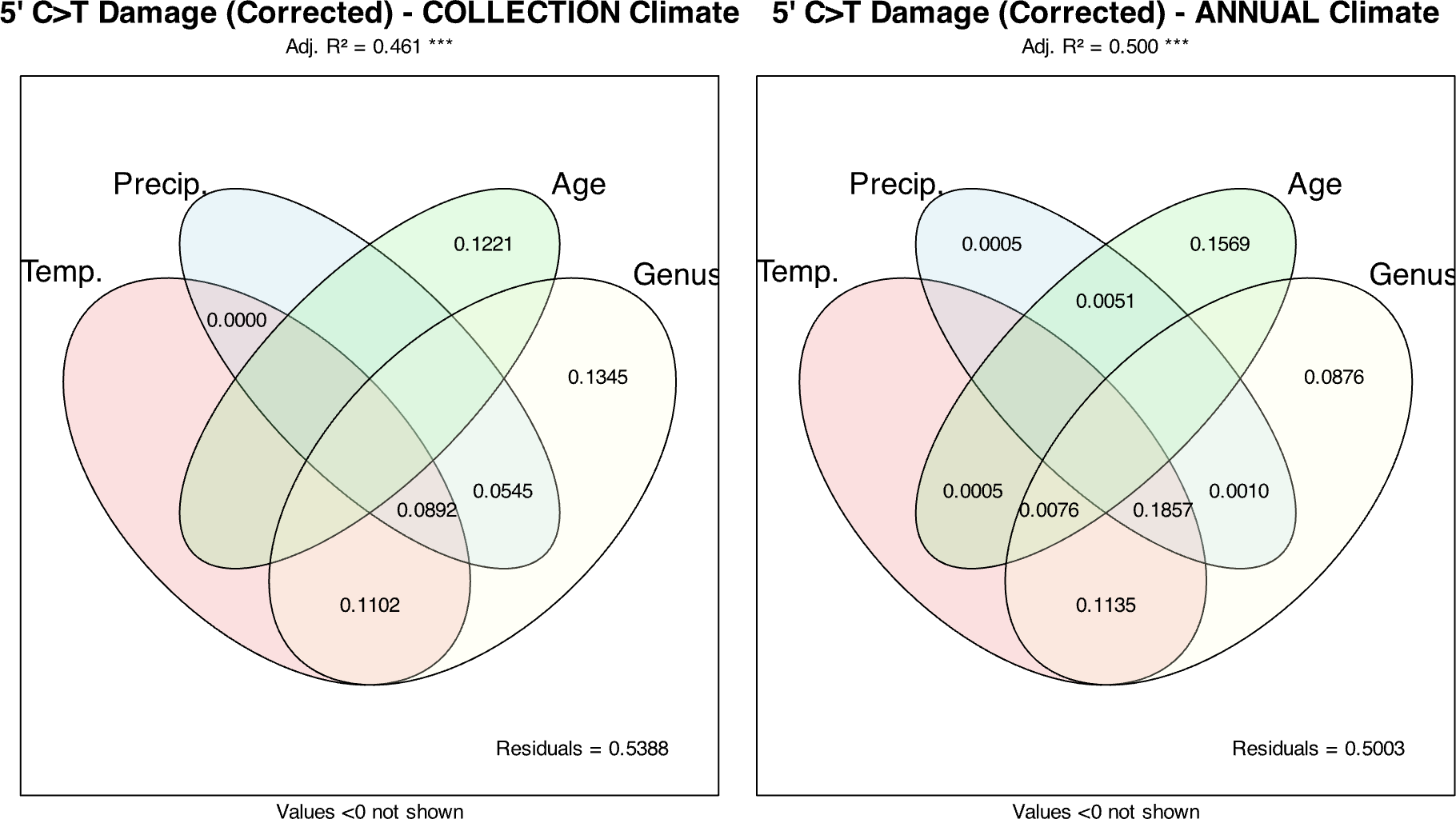

Supplementary figure S11: Variance partitioning analysis for 5’ C>T misincorporations frequencies corrected by baseline substitutions for the collection climate (left) and the annual climate models (right). Venn diagrams display the unique and shared contributions of the explanatory variables (temperature, precipitation, age and genus) to the total variance in corrected 5’ C>T misincorporations. Adjusted *R²* values for each model are shown above the plots. All reported values are statistically significant.

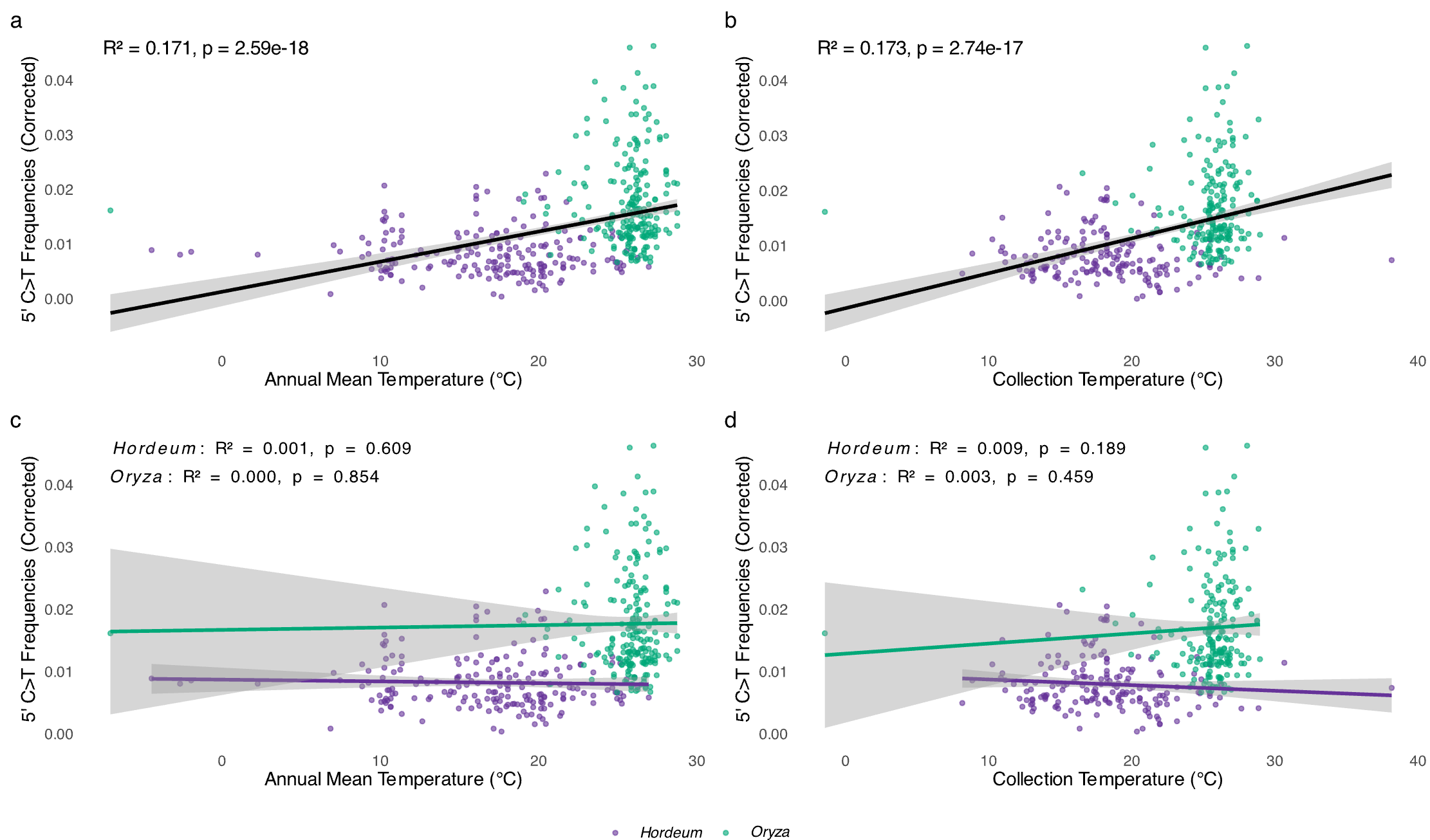

Supplementary figure S12: Relationship between temperature and corrected deamination rates (5’ C>T misincorporation frequencies corrected by baseline substitutions) in herbarium specimens for (a) annual mean temperature model, (b) collection temperature model, (c) annual mean temperature model for *Hordeum* and *Oryza*, and (d) collection temperature model for *Hordeum* and *Oryza*. Insets show regression statistics.
